# Depth-dependent eDNA abundances across ecosystems inform deep-sea sampling strategies

**DOI:** 10.64898/2026.05.12.724363

**Authors:** Santiago Herrera, Annette F. Govindarajan, Elizabeth Andruszkiewicz Allan, Rene Francolini, Erin Frates, Luke McCartin, Nicole C. Pittoors, Milan Sengthep, Sarah Stover, Samuel Vohsen, Nina Yang

## Abstract

Environmental DNA (eDNA) surveys are increasingly used to assess marine biodiversity and inform deep-sea environmental decision-making, including mineral resource management and fisheries oversight. Yet standard low-volume protocols inherited from coastal work may be inadequate at depth, and recent findings indicate that depth- and productivity-aware sampling, often requiring high-volume in situ filtration, is needed, but no quantitative framework links depth and ecosystem context to defensible filtration volume targets. We compiled 841 eDNA samples from eight expeditions across the North Atlantic, Wider Caribbean, and Pacific (surface to 4000 m) to quantify how recoverable eDNA scales with depth and surface productivity, and to derive depth- and productivity-aware sampling targets. Total eDNA concentration declined with depth as a power law, with attenuation exponents (*b*) modulated by surface productivity: most gradual in eutrophic waters (*b* = 0.67), intermediate in mesotrophic (*b* = 0.90), and steepest in oligotrophic systems (*b* = 1.25); volume-weighted models explained 66–88% of the variance. At a fixed extract-concentration target, required filtration volumes diverged ∼7-fold between oligotrophic and eutrophic systems at 200 m and ∼38-fold at 4000 m. Conventional Niskin sampling therefore undersamples deep-sea biodiversity, particularly in mid-to low-productivity systems. Among laboratory parameters, the assay-specific extract-concentration target exerted greater leverage on required volume than extraction efficiency or elution volume. Volume-aware sampling paired with optimized recovery should be routine in deep-sea eDNA surveys.

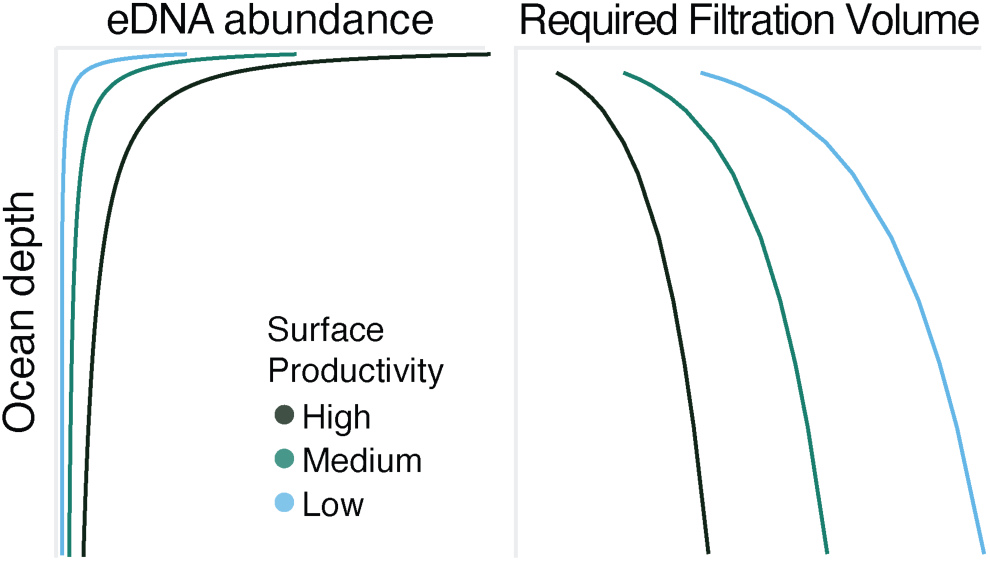

## INTRODUCTION

Aquatic environmental DNA (eDNA) surveys typically involve filtering water and extracting DNA from the resulting filter. The volume of water filtered is a critical determinant of detection sensitivity and inferred biodiversity. Yet in marine studies, filtered volumes remain predominantly small (typically 1–12 L, designated here as ‘low-volume’ sampling to distinguish it from the ‘large-volume’ sampling of tens to thousands of liters possible via high-capacity Niskins or high-volume in situ filtration), relative to the vast spatial dimensions and patchy heterogeneity of marine ecosystems. This constraint is especially pronounced in the ocean’s interior. The deep sea is the largest environment on Earth, comprising more than 90% of the habitable space, but eDNA studies there remain limited ^1^. Low organismal abundance, extreme pressures, restricted sampling opportunities, and high logistical costs pose additional challenges for deep-sea eDNA-based biodiversity assessments ^2–4^.

Most deep-sea eDNA sampling strategies are inherited from coastal, low-volume protocols. Still, recent work shows that deep-sea (>200 m, including the water column and near-bottom environments) animal eDNA surveys benefit from filtering larger volumes of seawater. Total measurable eDNA concentration, the eDNA mass recovered per L of filtered seawater, declines with depth ^2,5^. In the northwest Atlantic, McClenaghan et al. ^3^ found that samples below 1400 m yielded less DNA than surface and mid-depth samples, and that 1.5 L recovered more DNA and sequence variants than 250 mL. In the southern United States, Govindarajan et al. ^2^ showed that autonomous in situ sampler filtering ∼40–60 L detected ∼66% more metazoan taxa than ∼2 L Niskin samples; they also reported that the metazoan fraction of the eDNA signal declined with depth, implying that larger volumes become increasingly important in mesopelagic and deeper waters. Using deep-sea water in Japan, Yoshida et al. ^4^ concluded that 10– 20 L per replicate and >10 replicates were needed to reliably detect common fish species. Peres et al. ^6^ similarly showed that 1 L was insufficient to characterize deep-sea pelagic fish communities, whereas 5–10 L performed substantially better, and 5 replicates approached saturation. Together, these studies demonstrate the need for depth- and productivity-optimized eDNA sampling strategies.

Marine animal eDNA is also heterogeneously distributed and is insufficiently sampled with standard approaches, as metabarcoding replicates typically detect different taxa. Bessey et al. ^7^ found that aliquots of identical volume from the same tropical sample shared fewer than 60% of detected fish taxa, and Stauffer et al. ^8^ reported strong turnover between paired 30 L filtration replicates on tropical reefs, with assemblages differing in composition rather than forming nested subsets. Low-volume sampling, even at the surface, can therefore potentially produce incomplete and biased community assessments.

Post-sampling choices, such as extraction protocol, PCR replication, and sequencing depth, can improve biodiversity recovery, but their effects should not be conflated with sample-volume effects. Extraction efficiency varies among protocols ^9–11^; PCR replication increases detection ^6,12^; and sequencing depth, ideally sufficient for ASV rarefaction curves to plateau, is also important. In deep-sea environments where animal eDNA targets are rare, both sampling and post-sampling procedures should be maximized.

Despite these established observations, no quantitative framework currently exists to translate depth and ecosystem context into defensible filtration volume targets. This gap has practical consequences as eDNA evidence increasingly enters regulatory and policy contexts ^1^. This includes decisions made under the recently signed BBNJ Agreement (United Nations Convention on the Law of the Sea on the Conservation and Sustainable Use of Marine Biological Diversity of Areas beyond National Jurisdiction), deep-sea mineral resource management, marine protected area design, and fisheries monitoring ^1,13,14^. For these applications, the reliability of eDNA-based biodiversity assessments depends in large part on whether the underlying sampling was adequate to be representative of the targets of interest.

We address the fundamental operational question of how much seawater to filter as a function of depth by evaluating (1) how eDNA concentration scales with depth across different oceanographic settings, and (2) how filtration volume and post-sampling recovery contribute to biodiversity detection. We compiled measured eDNA concentration datasets from eight oceanographic expeditions across the North Atlantic, Wider Caribbean, and North Pacific, spanning oligotrophic, mesotrophic, and seasonally eutrophic ecosystems, and tested two hypotheses: (1) measurable eDNA concentration declines predictably with depth across ecosystems; and (2) under realistic operational ranges, increasing sample volume contributes more to the total eDNA mass recovered, that is, the total quantity of DNA captured on the filter summed across all organismal sources, than improving post-sampling recovery does. We use ‘total eDNA’ throughout to mean this bulk, fluorometrically quantified pool, dominated by microbial and other non-target sources, as distinct from the trace fraction attributable to any specific target taxon. By modeling eDNA concentration as a function of depth and distinguishing pre-from post-sampling considerations, this work provides regime- and depth-stratified filtration volume targets that researchers and practitioners can apply directly in survey design, including in contexts where eDNA evidence informs environmental decision-making.

## METHODS

### Field sampling

We compiled 841 environmental DNA (eDNA) samples collected over eight oceanographic expeditions between 2019 and 2023 in the eastern and western North Atlantic (*AR75*, *SG2105*), Wider Caribbean (*NF2202* and *FKt230417* off Puerto Rico; *AL2019* off the Bahamas; *MT19* and *PS2204* off the southern United States), and central Pacific near Hawaii (*NA155*; **Table 1**, **Figs. S1 and S2**; per-sample metadata in **Table S1**). Sampling depths ranged from the surface to 4000 m, with most samples from the epipelagic (<200 m) and mesopelagic (200–1000 m) zones. Two sampling approaches were used: (i) Niskin bottle rosettes; and (ii) Niskin bottles mounted on Remotely Operated Vehicles (ROVs) or Human Occupied Vehicles (HOVs) for near-bottom comparative samples on *FKt230417* (ROV *SuBastian*), *AL2019* (HOV *Nadir*), and *PS2204* and *NF2202* (ROV *Global Explorer*).

**Table 1.**
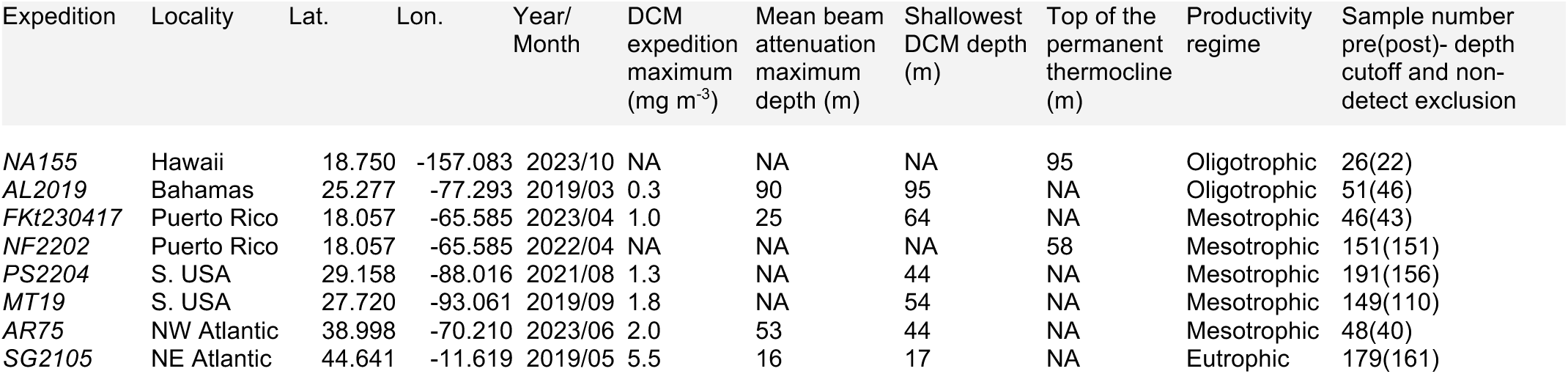
Characteristics of the eight oceanographic expeditions where eDNA samples were collected. The table details the locality, collection dates, deep chlorophyll maximum (DCM), mean beam attenuation, thermocline depth, productivity regime classification (oligotrophic, mesotrophic, or eutrophic), and the number of eDNA samples collected before and after filtering. Post-filtering counts are the samples remaining after the expedition-specific depth cutoffs and the further exclusion of samples at or below the assay limit of quantification.

Filtration used 0.22 µm polyethersulfone (PES) membranes throughout. Niskin-collected seawater (1– 10 L) was filtered shipboard through Sterivex cartridges or Smith Root capsules with peristaltic pumps ^5,15,16^. Filtered volumes were recorded for each sample (**Table S1**), and filters were stored at −80 °C.

### Total eDNA quantification

All filters, except those from *FKt230417,* were extracted with Qiagen DNeasy Blood & Tissue kits using format-specific modifications for Sterivex housings ^2,5,15^ and eluted in AE (Qiagen) or TE buffer. *FKt230417* filters were extracted by phenol/chloroform ^16^. We include this expedition because it is the only one for which paired target-eDNA (28S qPCR) measurements are available on the same extracts used for total-eDNA quantification, providing the empirical basis for the total-to-target relationships presented below, although it was excluded from productivity-stratified and mixed-effects modeling to avoid potential chemistry-based biases. Total eDNA was quantified using a Qubit fluorometer (Thermo Fisher), with the Broad Range (BR) assay and switching to the High-Sensitivity assay when concentrations were below the BR limit of quantification (LOQ). Template volume for quantification was 2 µL, and standards were run with every Qubit use. Final concentrations were standardized to ng L^−1^ of filtered seawater.

Because in four expeditions we performed both rosette- and vehicle-mounted (ROV/HOV) Niskin bottle sampling, we tested for systematic differences in eDNA concentrations between approaches using depth-controlled regressions within matched environmental strata, fitted with and without filtered volume as a covariate (Supporting Information, **Fig. S3**). No consistent effect of the sampling approach was detected, therefore samples from both approaches were retained in subsequent analyses.

### Modeling of eDNA concentration with depth

To describe how eDNA abundance changed with depth, we modeled the relationship using a normalized power-law function in R v4.3.1, adapting the Martin curve approach for vertical flux ^17^:

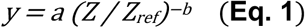

Here, *y* is the eDNA concentration (in ng L^−1^), *Z* is depth (in m), and *Z_ref_* is an expedition-specific reference depth used to normalize the depth axis across expeditions with different sampling ranges and productivity contexts. *Z^ref^* was set to the shallowest depth retained for each expedition after expedition-specific cutoffs that excluded observations above (in priority order): the mean transmissometer beam-attenuation maximum, the shallowest fluorometer chlorophyll maximum, or the top of the permanent thermocline, depending on data availability (**Table S1**; *Z_ref_* = 16–95 m). These features were strongly correlated with eDNA concentration (**Fig. S4**) and serve as practical proxies for the upper-ocean particle and chlorophyll maxima. These upper-depth cutoffs exclude near-surface observations shallower than the productivity-associated feature used to set *Z_ref_*, so that the model describes attenuation below the productive surface layer rather than within it. The normalization places each expedition’s cutoff at *Z*/*Z_ref_* = 1 and makes the attenuation exponent *b* comparable across expeditions.

We fit **Eq. 1** as a Gamma generalized linear model (GLM) with log link:

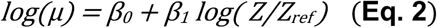

which is equivalent to the power law after back-transformation, *a = e*^*β*0^ and *b = −β_1_*. For each fit, we recorded *a*, *b*, 95% confidence intervals, the parameter variance–covariance, and pseudo-*R*² (1 − residual/null deviance). Models were fitted (i) globally, (ii) by surface productivity regime, classified from each expedition’s deep chlorophyll maximum (DCM) as oligotrophic (<1 mg m⁻³), mesotrophic (1–2 mg m⁻³), or eutrophic (>2 mg m⁻³), and (iii) by expedition. To reduce stochasticity and account for unequal filtered volumes, observations were also aggregated into normalized depth bins of width 0.5*·Z*_ref_; volume-weighted average concentrations (Σ ng DNA / Σ filtered volume) were then modeled per bin, excluding bins with ≤5 L total volume. Both unweighted and volume-weighted GLMs were fit to the binned data. The collective productivity effect was tested with a linear mixed-effects model of log-concentration on log(*Z*/*Z*_ref_) × productivity (fixed effects), with random intercepts and slopes by expedition and bin-volume weights. Significance was assessed by likelihood ratio test (LRT, df = 4) against a productivity-free reduced model, both refitted by maximum likelihood for fixed-effect comparison.

After cutoffs, 756 samples remained, of which 27 had non-detectable total eDNA (Qubit reading at or below the assay’s limit of quantification, LOQ; **Table S2**) and were excluded, leaving the 729 samples used in the fits above (**Table 1**). *FKt230417* was further excluded from the productivity-stratified and mixed-effects fits, to prevent confounding the extraction protocol with the productivity regime, because it used a different extraction chemistry; its per-expedition fit is reported separately.

Because non-detects were not distributed at random with depth, we tested the sensitivity of *b* to their exclusion. For each non-detect we reconstructed a sample-specific limit of quantification, LOQ_s_, in ng L⁻¹ (the assay’s LOQ in ng µL⁻¹ of extract scaled by elution volume and filtered volume), and refit the model three ways: substitution at LOQ_s_/2, substitution at LOQ_s_/√2, and left-censored (Tobit) regression of log-concentration on log(*Z*/*Z*_ref_) fit by maximum likelihood, with each non-detect treated as left-censored at its LOQ_s_. Because the censored model is Gaussian on the log scale, each treatment was compared against a complete-case Gaussian fit rather than against the Gamma GLM of **Eq. 2**. Re-including non-detect samples shifted *b* slightly upward, toward a steeper decline: by 0.11 in the oligotrophic and 0.10 in the mesotrophic regime (the eutrophic regime contained no non-detects), and by 0.04 in the pooled fit, in every case smaller than the corresponding confidence interval (**Table S3, Fig. S5**). Complete-case exclusion therefore did not inflate the depth-attenuation estimates and, if anything, rendered them slightly conservative.

### Calculating sampling volumes with depth

The seawater volumes required to recover a desired quantity of eDNA across depth were estimated by combining the fitted depth-attenuation models with post-sampling procedures. Because the upstream models were fitted on normalized depth, projection onto absolute depth requires an assumed reference. We set *Z_ref_assumed_* = 50 m for all regimes (representative of the fitted *Z_ref_* range, 16–95 m) so that the seawater volume required to recover a target total extract concentration *V(Z)* is directly comparable across regimes at any absolute depth. For each fitted model, expected total eDNA concentration at depth *Z* was calculated from the power-law relationship:

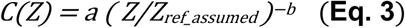

where *C(Z)* is the predicted eDNA concentration in ng/L. The mass of captured eDNA (*M_C_*) was defined as the quantity required to achieve a specified extract concentration after elution:

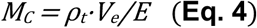

where *ρ_t_* is the total eDNA extracted concentration in ng L^−1^, *V_e_* is the final elution volume in µL, and *E* is the extraction efficiency.

The required filtration volume can be calculated as *V(Z) = M_C_/C(Z),* or replacing terms:

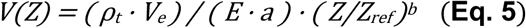

Baseline parameters were *E* = 0.80, within the upper range reported for Qiagen spin-column kits (0.3– 0.95; ^18,19^); *V_e_* = 80 µL, typical of our laboratory protocols ^2,15^; and *ρ_t_* = 8.1 ng µL⁻¹. Selecting a defensible value for *ρ_t_* is less straightforward because the minimum total eDNA extract concentration needed for reliable downstream detection depends on the specific assay, primer set, and target community. Because no general benchmark exists for *ρ_t_*, we derived an empirical worked-example value: the minimum extract concentration at which a 5 µL PCR aliquot would yield ≥20 copies (***ρ_min_***) of the 95th-percentile-rare *28S* amplicon sequence variant (ASV) observed in the *FKt230417* cnidarian/sponge/ctenophore community (^20,21^; full derivation including the qPCR assay and Zipf rank– abundance model in the **Supporting Information**). Reliable PCR amplification typically requires >10 target copies per reaction ^22^. We emphasize that this *ρ_t_* value is intended as a worked example rather than a universal recommendation. Practitioners applying this framework to other assays or target taxa should choose an appropriate *ρ_t_* for their own detection requirements.

Parameter uncertainty was propagated by Monte Carlo simulation. For each model, 5000 paired (log *a*, *b*) draws were taken from a bivariate normal parameterized by the GLM variance–covariance matrix; where only marginal 95% confidence intervals (CIs) were available, marginal standard deviations were derived from the CI half-width with zero covariance assumed. *C(Z)* and *V(Z)* were evaluated on a 200– 11000 m grid (predictions beyond 4000 m are extrapolations outside the empirical sampling range), retaining the median and the 5th, 25th, 75th, and 95th percentiles. Predictions were generated separately for the productivity-specific raw and volume-weighted bin models. Sensitivity of *V(Z)* to the operational parameters was assessed by sweeping *ρ_t_* (0.5–20 ng µL⁻¹), *V_e_* (20–150 µL), and *E* (0.2–1.0) one at a time across the prediction grid, and as a full factorial at depths from 200 to 4000 m, using the same cached parameter draws across scenarios to isolate parameter from sampling variance. This systematic approach allowed us to quantify how post-sampling procedures, such as adjusting elution volumes or optimizing extraction recovery, ultimately influence the field effort required.

Analytical code is available at https://github.com/herreralab/edna-depth-sampling-pipeline

## RESULTS

### Total eDNA concentration declines predictably with depth

Across all expedition datasets we examined, the total eDNA recovered per liter of seawater dropped sharply through the upper water column and then remained low at depth, despite the wide range of regions and seasons sampled (**Fig. S2)**. This pattern was well described by a simple power-law attenuation model (**Fig. 1, Table S4**). The fitted depth-attenuation exponent *b* was positive in all 27 fits across aggregation levels (expeditions and productivity regimes), with 95% CIs excluding zero throughout, confirming model adequacy and a consistent effect direction. Model fit improved markedly after depth binning and volume weighting: at the productivity-class level, pseudo-R² rose from 0.44–0.53 (raw data) to 0.66–0.88 (volume-weighted bins), with a median gain of 0.31 (**Table S4**). The volume-weighted bin model accounted for between 66% (eutrophic) and 88% (mesotrophic) of the variance in concentration with depth (**Table S4**). The practical implication is that depth is the dominant axis of variation in recoverable total eDNA concentration once expedition- and bin-level sampling variability is averaged out.

**Figure 1.**
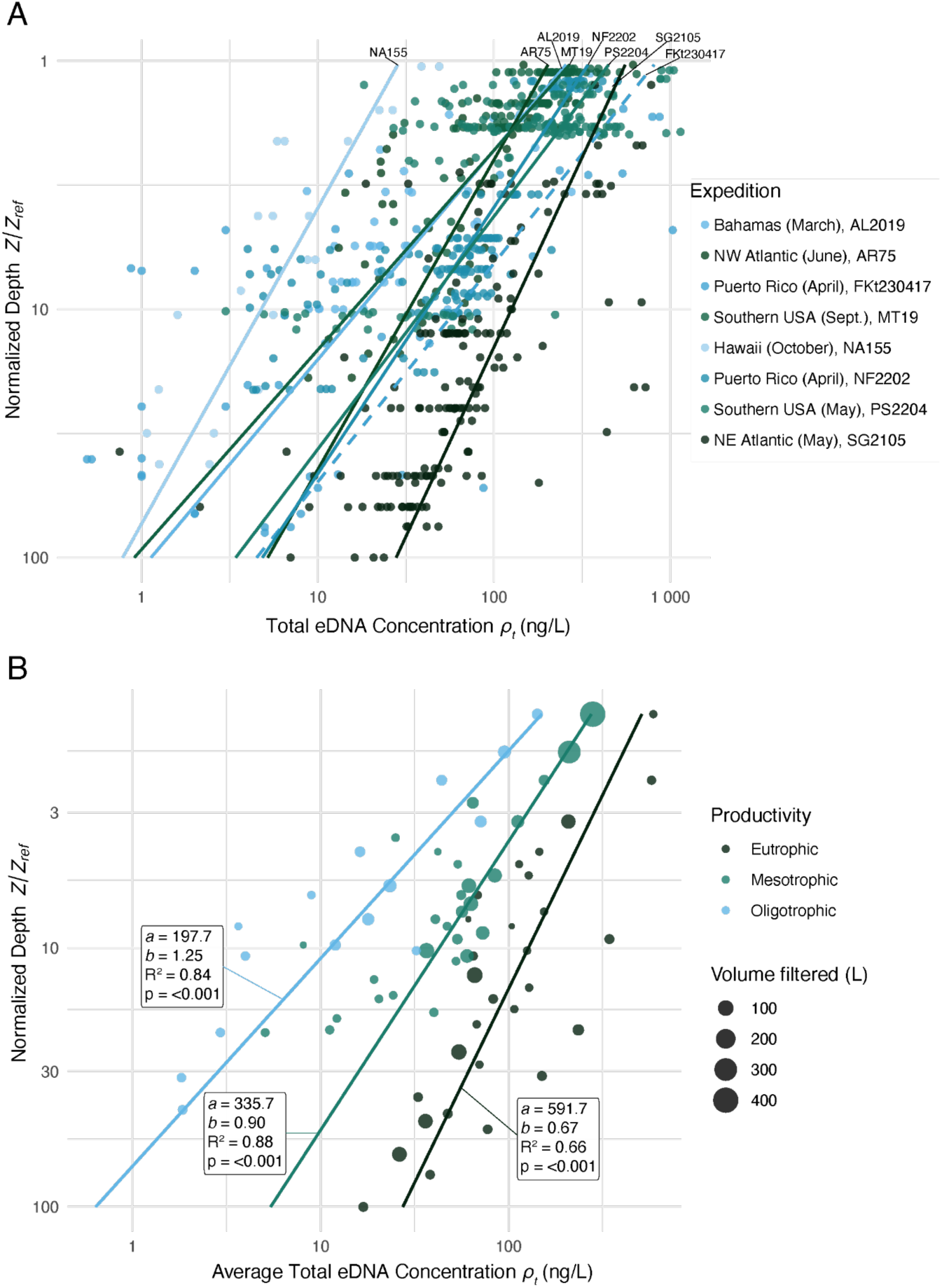
Vertical profiles of total eDNA concentration across depth, expeditions (A), and ocean productivity regimes (B). Panel (A) shows the raw observations colored by expedition. Panel (B) shows the volume-weighted binned averages colored by productivity classification, with point diameters scaled to aggregated filtered seawater volume per bin. Both axes are shown on a log scale. Solid lines represent the productivity-specific power-law models fitted using GLM with a Gamma error distribution and log-link function. The dashed line in panel (A) represents the fitted model for expedition *FKt230417* (phenol/chloroform extraction).

### Water-column productivity influences total measurable eDNA concentration

Expeditions in productive ecosystems consistently recovered more total eDNA at depth than expeditions in less productive ones. The near-surface concentrations and depth attenuation differed. Productivity regime was a strongly significant source of among-expedition variation in our dataset (likelihood-ratio test *p* = 1.5 × 10⁻⁵; *FKt230417* was excluded from this test because it used a different extraction chemistry).

Stratifying by productivity regime produced systematically different attenuation exponents (**Fig. 1, Table S4**): oligotrophic waters showed the steepest decline (*b* = 1.25, 95% CI 0.96–1.53; 84% of variance explained), mesotrophic waters were intermediate (*b* = 0.90, 0.79–1.00; 88%), and eutrophic waters the most gradual (*b* = 0.67, 0.44–0.90; 66%). This ranking held across raw and binned, weighted and unweighted fits, and explained considerably more variance than a single global fit (*b* = 0.47; 22%) (**Table S4**). Surface concentrations (intercept *a*) varied roughly threefold across regimes but with pairwise overlapping CIs. In contrast, *b* discriminated cleanly between the oligotrophic and eutrophic extremes — consistent with productivity acting more strongly on the rate of vertical attenuation than on the size of the upper-water-column reservoir.

The eutrophic regime is represented by a single expedition (*SG2105*) and should therefore be treated as illustrative of a seasonally productive system rather than as a generalized eutrophic estimate; the more robust contrast in our data is between the oligotrophic and mesotrophic regimes, both of which integrate multiple expeditions.

### Sampling volume requirements increase rapidly with depth

The sampling volume *V(Z)* required to meet the desired extracted eDNA yield in our worked-example (*ρ_t_* = 8.1 ng µL⁻¹, *V_e_* = 80 µL, *E =* 0.8, *Z_ref_assumed_* = 50 m) increased rapidly with depth (**Fig. 2**). For example, at 1500 m, our framework predicts that roughly 13 L of eutrophic waters, ∼51 L of mesotrophic waters, and ∼284 L of oligotrophic waters need to be filtered to recover the same amount of total eDNA. By 4000 m, those numbers rise to roughly 26 L, 123 L, and 966 L, respectively (**Fig. 2**). These are the volumes required to meet our *28S* worked-example extract-concentration goal; reliably detecting other target eDNA from that extract depends on that target abundance and assay-specific sensitivity (see **Discussion** and **Supporting Information**).

**Figure 2.**
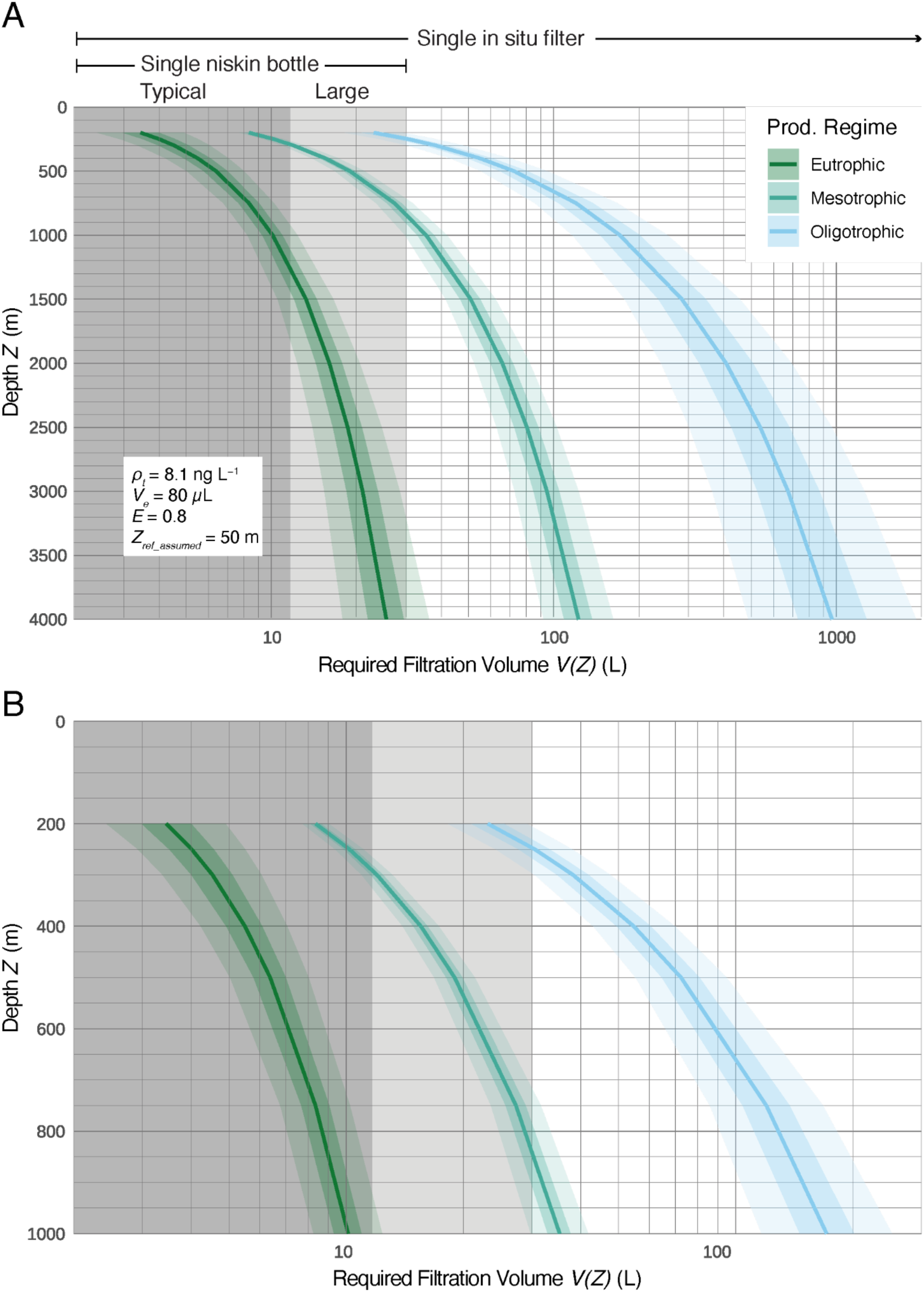
Predicted seawater volume, *V(Z),* required to achieve a specific eDNA concentration, *ρ_t_*, across depth and ocean productivity regimes. Panel (A) shows volume predictions in the deep sea between 200 and 4000 m. Panel (B) zooms in on the predictions in the mesopelagic, between 200 and 1000 m. The median required sample volume (L) is plotted against absolute depth (m). Colored lines represent the median predicted volume for each productivity regime, derived from a weighted, binned GLM with a Gamma error distribution and a log-link function. Shaded ribbons indicate uncertainty propagated by Monte Carlo simulation from the fitted parameter variance– covariance matrix, with the darker band showing the 50% interval and the lighter band showing the 90% interval. The x-axis is shown on a log scale. Shaded bands denote the approximate volumes of Niskin-based collection: ‘typical (small)’ Niskin bottles (about 1 to 12 L) and ‘large’ Niskin bottles (about 20 to 30 L). Predicted volumes above these bands cannot be met by a single bottle and imply either combining casts, at the cost of replication, or high-volume in situ filtration.

Because eDNA concentration declines with depth as a power law, the volume required to achieve any given extract-concentration target rises nonlinearly with depth, and the gap between productivity regimes widens accordingly: the predicted volume in oligotrophic versus eutrophic systems differs by ∼7-fold at 200 m, but by ∼38-fold at 4000 m. Deep-sea sampling effort is therefore especially sensitive to surface productivity. The main difference between sampling for eDNA in an oligotrophic ecosystem and in a productive ecosystem is the need to filter hundreds of liters rather than tens.

Uncertainty in these predictions also increases with depth and is largest in the oligotrophic regime. The 5–95% prediction interval in oligotrophic waters widens from ∼1.6-fold at 200 m to ∼3.9-fold at 4000 m (489–1914 L at 4000 m), whereas mesotrophic predictions stay within 1.2- to 1.8-fold and eutrophic predictions within ∼2-fold over the same range. The narrow eutrophic interval reflects only within-expedition variability, since a single expedition represents eutrophic; mesotrophic predictions are likely the most generalizable, as the mesotrophic fit integrates more samples across more expeditions (n = 5). Predictions from the raw-data model and the volume-weighted bin model agreed to within ∼2% in eutrophic waters and within ∼14% in mesotrophic waters across depth. Still, they diverged in oligotrophic waters at depth (by ∼19% at 1500 m and ∼24% at 4000 m, with the binned model predicting larger volumes), consistent with the steeper attenuation in those low-productivity waters.

### Scaling of operational parameters

Because *ρ_t_*, *V_e_*, and *E* enter *V(Z)* only through the multiplicative constant *M_C_* = *ρ_t_·V_e_/E* (**Eq. 5**), each rescales *V(Z)* by the same factor at every depth and in every productivity class, a property of the model, not an empirical finding. Changing any of these parameters, therefore, shifts the volume-versus-depth curve up or down without changing its shape (**Fig. 3**). Across the parameter ranges plausibly available to a practitioner, the target extract concentration *ρ_t_* has by far the largest leverage on required filtration volume. For example, reducing the target extract concentration *ρ_t_* by half requires a corresponding 50% reduction in the sampling volume. This multiplicative relationship extends to the other operational variables, extraction efficiency (*E*) and total elution volume (*V_e_*).

**Figure 3.**
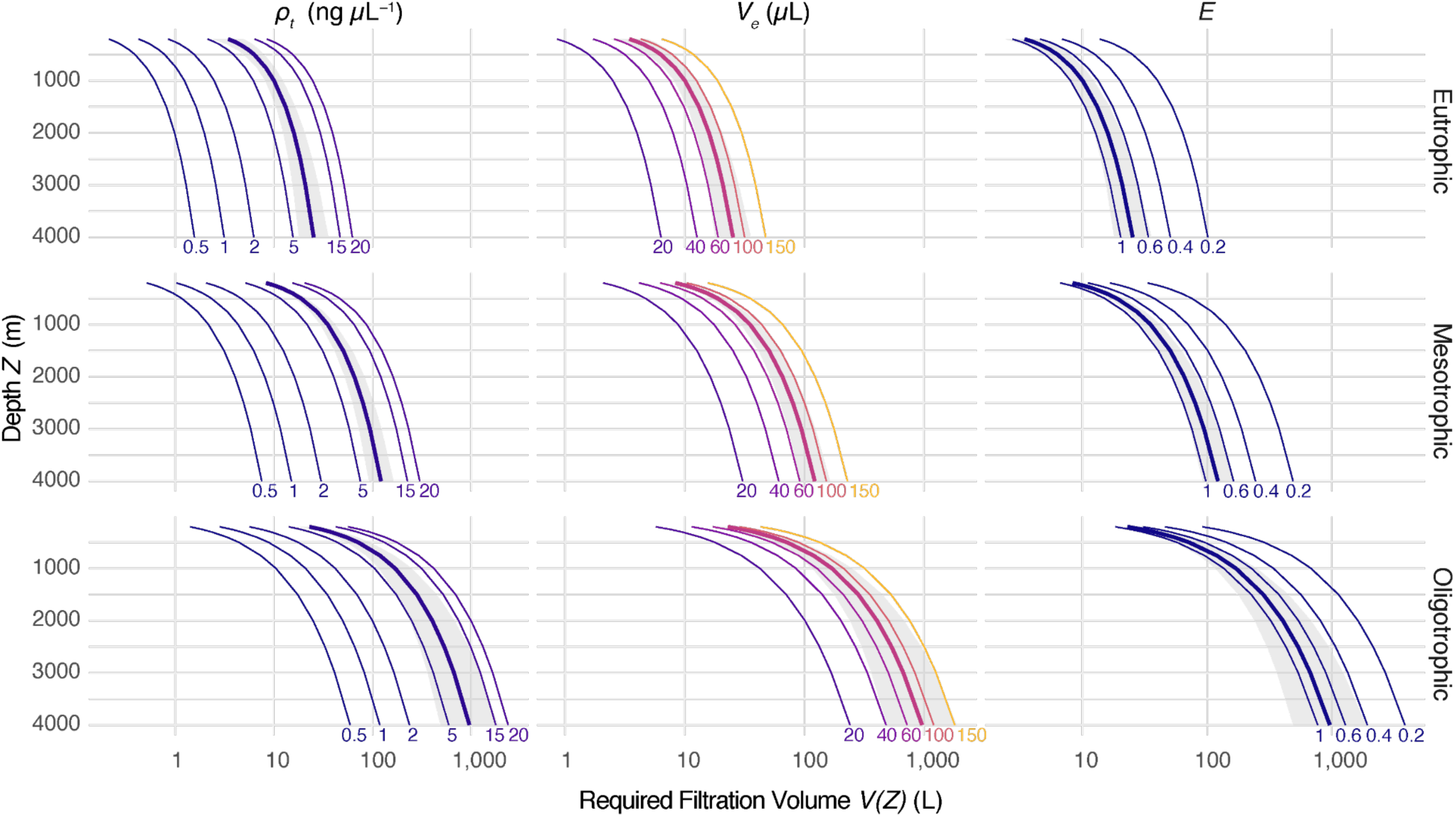
Sensitivity of required filtration volume *V(z)* to operational parameters (target extract concentration *ρ_t_*, elution volume *V_e_*, extraction efficiency *E*), predicted from the volume-weighted normalized-depth bin model by productivity regime (rows). Within each panel, the parameter under test is swept across the values labeled at the deepest point while the other two are held at baseline (*ρ_t_* = 8.1 ng µL⁻¹, *V_e_* = 80 µL, *E =* 0.8). The baseline scenario is drawn with a thicker line, and its 5–95 % Monte Carlo band is shown in grey. All predictions assume *Z_ref_assumed_* = 50 m.

Across the parameter ranges tested, *ρ_t_* varied *V(Z)* by 40-fold (0.5–20 ng µL⁻¹), *V_e_* by 7.5-fold (20–150 µL), and *E* by 5-fold (0.20–1.00). Combining all three produces a ∼1500-fold envelope (0.012× to 18.5× the baseline), which is constant across depths and regimes. The single most consequential operational parameter is therefore the assay-specific target concentration. Relaxing *ρ_t_* from 8.1 to 0.5 ng µL⁻¹ alone reduces required volumes ∼16-fold, more than can be achieved by improving extraction efficiency or reducing the elution volume within realistic laboratory ranges (**Table S5, Table S6, Fig. S6**).

For our worked example (*28S* amplicon assay, *ρ_t_* = 8.1 ng µL⁻¹, *V_e_* = 80 µL, *E =* 0.8, *Z_ref_assumed_* = 50 m), the required sample volumes spanned nearly two orders of magnitude across productivity regimes at any given depth. Practitioners with less stringent detection requirements (lower *ρ_t_*) or more efficient laboratory workflows (higher *E*, smaller *V_e_*) can scale these numbers down accordingly using **Eq. 5** and the regime-specific parameter estimates for *a* and *b* (**Fig. 2**).

## DISCUSSION

This study provides a quantitative framework for optimizing deep-sea eDNA sampling. Recoverable total eDNA concentrations decline predictably with depth as a power law across the oceanographic settings sampled here. Beyond depth, two factors define the sampling effort needed to recover usable amounts: surface productivity, which likely governs how rapidly eDNA attenuates with depth, and post-collection recovery, which determines how much of the captured material reaches downstream analyses. Below, we consider each factor, discuss limitations, and provide operational guidance for deep-sea eDNA sampling.

### Total eDNA concentrations predictably decline with depth

Below the productive upper-ocean layer, recoverable total eDNA decreases steadily with depth, analogous to the Martin curve for sinking particulate organic carbon ^17^. In both cases, particle-associated material is degraded, diluted, dispersed, and removed from suspension as it descends. This unifying interpretation aligns with empirical studies showing that increased seawater filtration volumes consistently improve DNA yields and taxonomic recovery in the deep sea ^2–4,6^. It explains why that pattern is robust across systems. The steep near-surface decline followed by persistently low concentrations at depth implies that recoverable eDNA in the deep sea is ubiquitously scarce relative to the volumes typical of coastal or freshwater work, and that scaling deep-sea sampling using those protocols is, in many cases, fundamentally inadequate.

### Productivity modulates eDNA depth attenuation

Surface productivity is an important ecological correlate of marine eDNA distributions. Greater standing biomass and stronger export flux in productive systems should enhance the downward supply of organic material and, therefore, associated DNA, resulting in a slower decline of eDNA concentration with depth. Oligotrophic waters, with reduced biomass and vertical flux, should attenuate more rapidly. Our observations and model fits are consistent with this expectation. Attenuation exponents (*b*) decreased systematically from oligotrophic to eutrophic, with the clearest separation between the oligotrophic and eutrophic regimes. The confidence intervals of fitted intercepts (*a*), in contrast, overlapped across regimes despite a threefold spread in point estimates. Together, these patterns suggest that productivity is more closely tied to the rate of vertical attenuation than to the magnitude of the upper-water-column eDNA pool. However, the weak separation may also reflect that our sampling did not specifically target subsurface chlorophyll or particle maxima.

Two caveats temper this interpretation. First, the eutrophic regime fit is based on a single expedition (*SG2105*); it should be regarded as illustrative rather than representative of the productivity regime. The more robust contrast, supported by our data, is between oligotrophic and mesotrophic regimes, both of which integrate multiple expeditions. Second, productivity regime classification was derived from expedition-level chlorophyll maxima and so confounds productivity with region, season, and other ecological contexts. Future studies that explicitly disentangle productivity from these covariates and that expand replication at the eutrophic end of the gradient would strengthen this interpretation.

### Uncertainty in sampling design for target volumes

Uncertainty in the required sample volume to achieve a desired eDNA quantity grows with depth. In the upper ocean, where modeled eDNA concentrations are higher, predicted filtration requirements remain comparatively constrained. As concentration declines in deeper water, uncertainty in the fitted relationship propagates into much broader volume estimates, with the greatest uncertainty in deep oligotrophic regimes. This uncertainty has direct implications for sampling design. Deep-sea eDNA surveys should adopt conservative sampling volumes, since underestimating filtration requirements will likely reduce detection probability and bias downstream comparisons. In practice, this underscores the need for in situ samplers with high flow rates that can filter large volumes in tractable time windows ^2,23,24^. In many deep-sea environments, Niskin bottle sampling cannot obtain sufficient volumes to adequately sample eDNA, and combining bottles into a single filter to reach larger sample volumes reduces replication and still does not approach the volumes that high-flow in situ samplers can filter. Our findings therefore support adaptive sampling designs that allocate more effort to deeper, less productive ecosystems where logistically feasible, and a broader push to make large-volume in situ sampling technologies accessible as deep-sea eDNA surveying expands.

### Field effort vs. post-collection recovery

Our results also imply that post-collection laboratory steps, such as DNA extraction, PCR replication, and sequencing depth, should be central components of deep-sea eDNA analysis pipelines. It is important to distinguish between efforts that increase biodiversity detection by collecting larger sample volumes versus those that focus on post-collection processing steps. Increasing the recovery effort cannot compensate for inadequate sampling volume because it only retrieves eDNA molecules captured in the original field sample; it does not recover eDNA fragments that were never collected.

Increasing the extraction-efficiency parameter *E* reduces the modeled required water volume by a constant proportional factor across all depths and productivity regimes (**Eq. 5**, **Fig. 3**). The largest absolute reductions therefore accrue in the deepest, least productive waters, where baseline predicted volumes are largest, even though the relative gain is the same everywhere. This pattern aligns with broader eDNA literature showing that extraction, inhibitor removal, and amplification choices strongly influence taxa detection and false-negative rates, especially when target eDNA abundances are low ^10,25–28^. It also aligns with the recent emphasis on fit-for-purpose protocol optimization rather than universal workflows, especially across environments that differ in turbidity, inhibitor load, and expected target eDNA abundance ^28,29^. Among the parameters that practitioners can adjust, the assay-specific target extract concentration *ρ_t_* has by far the largest leverage on required filtration volume, because its effect within the evaluated range (∼0.5–20 ng µL⁻¹, ∼40-fold) substantially exceeds the realistic ranges for elution volume *V_e_* and extraction efficiency *E*. Practitioners targeting relaxed detection thresholds (e.g., targeting a more abundant taxon) can therefore reduce required field sampling volumes more by selecting more sensitive downstream assays than by improving any single laboratory recovery step.

Improving lysis efficiency, extraction chemistry, or inhibitor mitigation can partially offset field effort by reducing volume requirements, shortening filtration times, and easing logistical burdens such as pump capacity, clogging, and platform limitations. These gains are hard to quantify, however: extraction efficiency must be measured by recovery experiments, and reported values span a wide range (∼0.3– 0.95; ^18,19^) reflecting both method differences and strong sample dependence. Inhibitors introduce a further complication that runs counter to the depth-attenuation gradient, as they accumulate alongside eDNA on the filter, so larger filtered volumes may also carry more inhibitors ^25,29^. In our experience, inhibition tends to be a bigger problem in productive ecosystems, whereas deep oligotrophic samples yield less DNA and cleaner extracts. The net effect on amplifiable recovery is therefore not straightforward and is not routinely measured or reported. Deep-sea eDNA studies should therefore consider improving the recovery of amplifiable target eDNA (e.g., by adding DNA clean-up steps to their laboratory protocols), which may be as consequential as increasing the volume of seawater collected. While deep-rated, large-volume in situ samplers are expensive and remain the exception rather than the rule in eDNA studies ^24^, improving post-recovery efficiency may be a broadly tractable approach.

Because extraction efficiency and inhibition are rarely measured yet strongly influence recovery, we further recommend that future deep-sea eDNA studies incorporate internal spike-in standards ^30,31^. These standards could be exogenous or synthetic DNA added at a known quantity prior to extraction, to estimate per-sample recovery efficiency and quantify amplification inhibition directly, which we expect to be more severe in productive surface waters than in deep oligotrophic samples.

Finally, a decline in total eDNA with depth does not by itself preclude biodiversity recovery. When sequencing depth is adequate for per-sample accumulation curves to plateau, common taxa can be characterized even from modest yields. What the decline sets is a floor to the minimum quantity of target template physically present in the sample that sequencing can detect. Within-sample ASV richness typically saturates near 10⁵ to 10⁶ reads per sample per marker ^32–34^ Beyond that sequencing adequacy threshold the constraint on rare-target recovery shifts from sequencing effort to the amount of template captured in the field. Detected richness, by contrast, rarely saturates with increasing filtered water volume. At a fixed marker and sequencing depth, observed richness scales with cumulative replicates/volume along log–log slopes of approximately +0.4 to +0.5 ^8,35^. The two controls are therefore distinct. Sequencing is no substitute for a template-sufficient field sample, which provides the quantitative rationale for prioritizing volume and replication in deep-sea eDNA survey design.

### From total eDNA to target detection

The predicted sampling volume required to achieve a desired total eDNA concentration should be treated as a minimum when targeting specific taxa. In most cases, eDNA from target animal taxa constitutes only a tiny fraction of total marine eDNA. For example, using a qPCR assay with the *28S* primers that specifically target cnidarians, ctenophores, and sponges ^20,21^, we estimated that our near-bottom samples from Puerto Rico (expedition *FKt230417*; 2 to 2.5 L per sample) contain between 0.00001% and 0.0001% of the total eDNA mass recovered from each extracted filter (**Table S7**). This range is consistent with estimates by Andruszkiewicz Allan et al. ^25^, and Stat et al. ^36^, where their target eDNA was only 0.00001% (targeting dolphins in captivity by qPCR) and 0.00004% (targeting fish in a coral reef by shotgun sequencing) of their total eDNA, respectively. Despite this small fractional contribution, total eDNA and *28S* target eDNA were strongly correlated across the 24 Niskin samples with quantifiable target signal (Pearson r = 0.95, p = 2×10^-12^; **Table S7**, **Fig. 4**). Within this single assay-and-expedition combination, total eDNA therefore tracks the *28S* target signal closely, supporting the practical use of total-eDNA volume calibrations as proxies for target-detection volumes within a given assay context. However, the generality of this correlation across primer sets, target taxa, productivity regimes, and expeditions remains to be tested.

**Figure 4.**
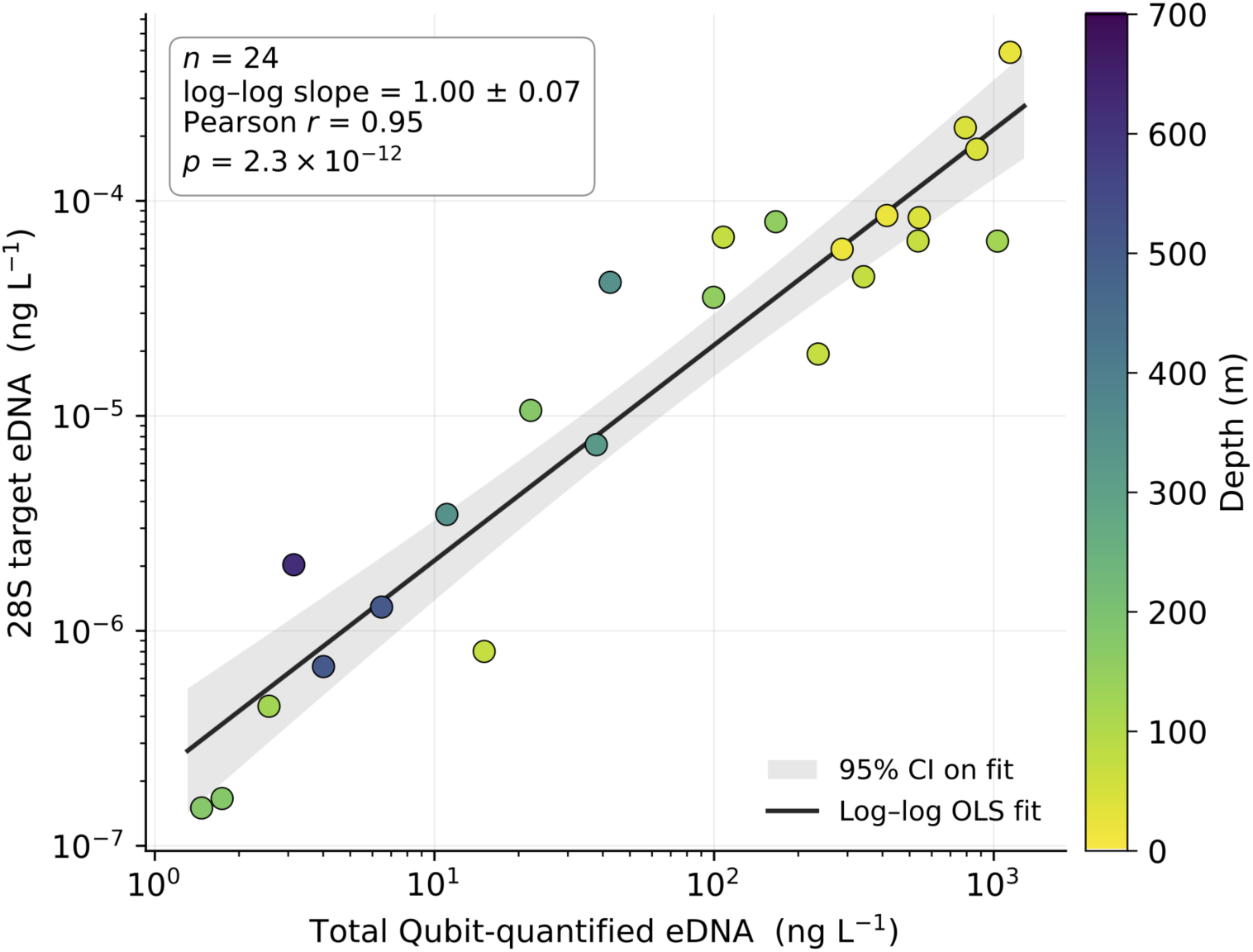
Total Qubit-quantified eDNA versus 28S-target eDNA across Niskin samples from the *FKt230417* expedition (Puerto Rico). Each point is one filter (*n* = 24; 0.22 µm Smith-Root capsules; 2.0–2.5 L filtered per sample), colored by sampling depth (16–616 m). The solid line is an ordinary least-squares fit in log space, and the grey ribbon shows its 95% confidence interval. Total and target eDNA scale proportionally across three orders of magnitude in total concentration (log-log slope = 1.00 ± 0.07; Pearson *r* = 0.95, *p* = 2 × 10⁻¹²), indicating that total Qubit-quantified eDNA is a good proxy for target-taxon eDNA within this single assay–expedition combination. The 28S qPCR primer set targets cnidarians, ctenophores, and sponges ^20,21^.

For species-specific assays such as qPCR, detection sensitivity depends not only on the eDNA mass captured on the filter but also on the fraction of the resulting extract that is actually queried per reaction. The absolute template volume, the template-to-reaction-volume ratio, and the number of technical replicates jointly set the effective limit of detection for rare targets ^6,12,26^, and our supplementary rank-abundance framework shows that template volume alone modulates required filtration volume by ∼10-fold across realistic ranges (1–10 µL; **Eqs. S6-S7, Fig. S7**). Expanding template volume or running additional replicates can therefore recover detections that small extracts would otherwise miss, but cannot rescue a field sample that was too small to contain the target in the first place.

Together, these estimates indicate that target deep-sea animal eDNA is typically found in trace amounts, equivalent to tens to hundreds of target eDNA molecules per liter. Therefore, adequate sampling of target eDNA from animal taxa must be proportionally greater than the sampling needed to characterize microbial biodiversity through DNA, which can make up the majority of the total eDNA (>60%) for a given sample ^36–38^. The framework presented in this paper should therefore be interpreted as an approximate predictor of recoverable DNA material, not as a direct predictor of the entire suite of biodiversity across markers or taxa, and not as an indicator of post-collection recovery inclusive of amplification and sequencing strategies.

### Caveats and assumptions

As discussed above, total eDNA is not a direct proxy for biodiversity. Two further considerations highlight this caveat for deep-sea applications. First, methods that maximize total eDNA do not always maximize target eDNA or the target-to-total ratio, especially when inhibitors or abundant off-target eDNA are co-concentrated ^25^. However, in our *FKt230417* Puerto Rico dataset, the two tracked closely (**Table S7**, **Fig. 4**). These filters were extracted by phenol/chloroform rather than the Qiagen chemistry used elsewhere; while the total-to-target relationship is consistent within this expedition, the absolute extract yields, and hence this ratio, may not transfer directly to Qiagen-based protocols. Different primers or taxonomic targets may respond differently to the same field design ^15^. Second, the metazoan fraction of total eDNA itself declines with depth ^2,5,39^, so the animal-specific attenuation exponent is likely steeper than the total-eDNA attenuation exponent reported here, implying that *V(Z)* for animal-targeted surveys at depth is likely larger than the volumes our framework predicts.

Our sampling did not target deep sound-scattering layers, where animal biomass and diversity ^40–42^ are concentrated and where eDNA accumulates locally. Also, for the subset of animals that undertake diel vertical migration – moving between mesopelagic and surface waters – the vast majority of their eDNA is predicted to be at the surface and mesopelagic depths where those animals spend most of their time^43^. This has been observed in the field ^44^ and in metabarcoding results demonstrating diel shifts in biodiversity ^45–47^. Similarly, in this study, our single AR75 day-night paired sampling cast suggested a surfaceward shift in eDNA at night (**Fig. S8**). Future studies should explicitly consider how to optimize sampling approaches, including incorporating targeted sampling, with respect to both migrating and non-migrating mesopelagic animal biomass ^44^.

The predictive framework also depends on several assumptions. Projected sample volumes rely on fixed parameter values for extraction efficiency, elution volume, desired recovered concentration, and the assumed reference depth. Our empirical dataset extended to 4000 m, with most collections limited to the top 1000 m, while volume predictions were generated to 11000 m. These deeper predictions illustrate how effort scales under the fitted model but remain extrapolations beyond the observed range. Likewise, using an expedition-normalized reference depth provides an analytically useful basis for comparison, but translating these relationships back to absolute depth in a new system depends on direct measurements or assumptions about the reference depth. In particular, eutrophic or nutrient-enriched systems typically have shallower DCMs than oligotrophic systems, where DCMs commonly occur much deeper in the water column, often near the deep nutricline ^48^. Thus, applying a single *Z_ref_assumed_* = 50 m projection likely compresses the cross-regime separation in *V(Z)* that would emerge if specific reference depths were used. Future work could evaluate these assumptions and extrapolations across seasons, basins, and target taxa, and their link to marker-specific richness saturation, occupancy-based detection goals, and other community-level metrics.

### Practical guidance for sampling design

The framework presented here incorporates depth, surface productivity, and laboratory parameters to predict filtration volumes *V(Z)* (**Eq. 5**), where *a* and *b* are the productivity-regime-specific power-law parameters (**Table S4**), *ρ_t_* is the target total extract concentration, *V_e_* is the elution volume, *E* is the extraction efficiency, and *Z_ref_* is the reference depth. At a fixed extract-concentration target, the predicted filtration volume diverges by roughly 7-fold between oligotrophic and eutrophic systems at 200 m and by ∼38-fold at 4000 m, meaning that a single ’deep-sea’ sampling protocol cannot serve all regimes. These cross-regime comparisons are evaluated at a common reference depth (*Z_ref_assumed_* = 50 m) so that regimes can be compared at equal normalized depth; they are not predictions for specific real-world sites, whose differing reference depths would likely widen the divergence.

When *ρ_t_* is unknown, or if there are multiple targets across a range of concentrations, we recommend using a conservative rarity-aware value as a starting point. As described above, the measured proportion of target eDNA from a single species or an animal single-clade out of total eDNA ranges between 10^−6^ to 10^−7^, but is likely to be higher in high-abundance situations such as feeding or spawning aggregations. These fractions imply a *ρ_t_* of 1 to 10 ng µL⁻¹, but they may be < 1 ng µL⁻¹ for very high-abundance taxa. When feasible, we recommend performing a qPCR/ddPCR quantification to adjust the per-reaction copy threshold ρ_min_ from the validated limit of detection or quantification of the chosen assay, and a sequencing pilot to obtain the target rank *n* from the target taxon’s abundance in the amplifiable community (absent prior information, a conservatively high rank, i.e. 90th to 95th percentile of expected richness) and the Zipf exponent *s*. Alternatively, *n* and *s* can be calculated from published environmental amplicon data or rank-abundance summaries for the same marker and depth zone.

Because *V(Z)* is formulated against total eDNA, it constitutes a minimum filtration volume. Detection of specific taxonomic targets, for example, fish, coral, or sponge species via metabarcoding or qPCR, requires the additional rescaling described in the **Supporting Information**, which links *V(Z)* to the rank-abundance distribution of the amplifiable target community via a truncated Zipf model. Practitioners applying this framework should therefore (i) select *ρ_t_*, *V_e_*, and *E* values appropriate to their laboratory workflow; (ii) substitute their own site-specific *Z_ref_*; and (iii) when targeting specific taxa, propagate *V(Z)* through the rank-abundance step, parameterized by the target ASV rank *n*, the community Zipf exponent *s*, and the per-PCR reaction copy threshold *ρ_min_*.

Because *Z_ref_* was set to each expedition’s empirical cutoff depth, applying *V(Z)* to a new system requires either a measured site-specific reference depth or an assumed value for *Z_ref_*. Practitioners can substitute their own reference depth (e.g., the deep chlorophyll maximum, transmissometer maximum, or top of the permanent thermocline) by rescaling the predicted volume by (*Z_ref, site_* / 50)*^b^*, where 50 m is the reference depth used in our worked predictions.

As approximate field-planning guidance based on the upper bound of the 90% uncertainty interval for total eDNA recovery, we suggest minimum filtration aims of single to tens of liters in eutrophic systems (achievable with typical low-volume 1-12 L, and large 20-30 L, Niskin bottles), tens to a few hundred liters in mesotrophic systems (achievable with large Niskin bottles and in situ filtration), and hundreds to thousands of liters in oligotrophic systems (achievable only with in situ filtration). These bounds should be adjusted to reflect the relative abundance of the target taxon.

Our predictions are based on samples filtered through 0.22 µm membranes, which is typically the smallest pore size used by eDNA practitioners. In our experience, small-area filters (e.g., 47 mm and 90 mm disk membranes; 17-64 cm^2^) tend to limit the filtered volume by reducing flow rates and can clog, but are easy to process post-sampling and are relatively economical (< $2 each). Large-area filters (e.g., Waterra filters; 300 cm^2^) enable high flow rates and rarely clog; pers. obs.), but the post-sampling processing burden is significant, and they are more expensive (Waterra filters are ∼$14 each, comparable to Sterivex). Larger pore sizes (e.g., 1.2 or 5 µm) allow the filtration of larger volumes at faster flow rates per unit of filter area, and generally capture a large proportion of target metazoan eDNA ^2,25,38^. However, animal taxa shedding smaller eDNA particles may be missed, as may microbial taxa. The trade-offs between filter pore size and area must be carefully evaluated to align with the sampling objectives.

Lastly, the optimal practical strategy varies according to whether the research focuses on targeted single-species detection (e.g., via qPCR or ddPCR) or broader community profiling (e.g., via metabarcoding). Because our volume models are parameterized on total eDNA, they establish a template-availability baseline applicable across both methodologies. However, for targeted qPCR or ddPCR, detection is primarily sensitive to the total mass of the target template and the fraction per reaction. In contrast, the completeness of metabarcoding-based assessments is further modulated by sequencing depth, primer-induced biases, and taxonomic richness saturation. While achieving a fixed total-eDNA concentration may meet the requirements for species-specific assays, community-level characterizations are often constrained by sequence rarefaction or replication; the filtration volumes derived here therefore constitute a baseline that must be paired with optimized recovery and sequencing effort to achieve representative biodiversity coverage.

### Implications for future studies

eDNA surveys are emerging as one of the most tractable tools for scaling biodiversity assessment in the deep ocean ^1^, and have become a routine component of deep-sea expeditions ^49^. Given urgent societal pressures from environmental change, fisheries harvesting, and mineral resource extraction, the depth- and productivity-dependence of eDNA recovery has direct implications for monitoring program design. Our results indicate that standard low-volume Niskin-bottle approaches will systematically underdetect animal biodiversity at depth, whether in the mesopelagic, where animal biomass is substantial ^41,50^, or at the deep seafloor, where species diversity can be high and biomass low ^51^. In situ sampling technologies capable of filtering the requisite tens to hundreds of liters within tractable time windows remain limited and are in the early stages of technological readiness ^24^. Passive samplers offer a complementary low-cost route to “large-volume” sampling. Although passive samplers do not actively filter water, integrating them into midwater trawls, towed bodies, or other gear that already passes very large volumes of seawater past the substrate effectively couples a passive medium to a high-throughput sampling stream ^52,53^. This pairing is worth further exploration as a near-term option, particularly in the midwater where plankton-net deployments are already routine.

While technology advances rapidly, we expect low-volume Niskin sampling to likely remain a common mode for deep-sea eDNA work in the near term. Substantial ecological insights have been and will continue to be gained from Niskin-based studies ^15,33,46,47,49,54^. Notably, Niskin-collected eDNA also detects more animal taxa than plankton nets despite nets passing far larger water volumes, a function of net mesh-size selectivity and sample-handling losses rather than of filtered volume per se ^5,23,33,55^. Our framework predicts that volume sets a detection floor for low-biomass or low-shedding community members, even when common taxa (e.g., microbes) can be recovered from low-volume samples.

Taken together, accounting for predictable depth-related declines in recoverable DNA, ecosystem productivity, and post-collection recovery efficiency should be a routine part of deep-sea eDNA sampling design. The framework outlined here identifies the depth–productivity combinations under which conventional low-volume sampling is most likely to produce biased coverage, and where additional investment in volume, replication, and analytical optimization will yield the largest gains. Adequate, representative sampling is especially important as eDNA surveys increasingly inform environmental decision-making in the deep sea.

## ASSOCIATED CONTENT

### Supporting Information

Supporting methods for the 28S qPCR assay, the truncated Zipf rank-abundance framework, the comparison of rosette- and vehicle-mounted sampling, and the diel and near-bottom observations; Figures S1 to S10 (PDF)

Per-sample eDNA quantifications and collection metadata, non-detect samples and reconstructed quantification limits, fitted depth-attenuation parameters, predicted filtration volumes, paired total and 28S-target quantifications, and Zipf rank-abundance summaries; Tables S1 to S11 (XLSX)

## ACKNOWLEDGEMENTS

We thank the crew and science teams on the expeditions *AR75* on the R/V *Neil Armstrong* (Chief Scientist: Andone Lavery), *SG2105* on the R/V *Sarmiento* de *Gamboa* (Chief Scientists: Ken Buesseler and Heidi Sosik), *MT19* on the R/V *Manta* (Chief Scientist: Santiago Herrera), *PS2204* on the R/V *Point Sur* (Chief Scientist: Santiago Herrera), *NF2202* on the NOAA Ship *Nancy Foster* (Chief Scientist: Andrea Quattrini), *FKt230417* on the R/V *Falkor (too)* (Chief Scientist: Colleen Hansel), *AL2019* on the R/V *Alucia* (Chief Scientists: Heidi Sosik and Joel Llopiz), and *NA155* on the E/V *Nautilus* (Expedition Lead: Jason Fahy, Science Leads: Larry Mayer, Chris Roman, Annette Govindarajan). We thank C. Park (WHOI) for her assistance with the illustration in the graphical abstract.

## SUPPORTING INFORMATION

### 1. Diel vertical migration and benthic boundary observations

Ecosystem- and depth-specific phenomena may produce pulses of animal biomass and biodiversity that affect the relationships among total eDNA yield, target-specific eDNA concentrations, and depth. Specifically in the top 1000 m, the presence of deep sound-scattering layers, some of which undergo daily vertical migrations, may cause fluctuations in eDNA relative to constituent biomass ^1,2^. Near-bottom environments, likely including a mixture of water-column and benthic eDNA, may also affect the relationship between eDNA yield and depth. While our sample sets were insufficient to assess these phenomena thoroughly, we conducted a first-order analysis for expeditions when the sample sets were amenable.

To assess whether total eDNA yields measurably fluctuate due to diel vertical migration, we compared eDNA yields from the *AR75* expedition in the Northwest Atlantic. The *AR75* was the only dataset available that included high vertical resolution and back-to-back paired daytime and nighttime casts. To assess whether proximity to the seafloor influenced eDNA samples, we plotted eDNA yields against altitude (height above bottom) for expedition *PS2204*, which included near-bottom samples (**Fig. S9**).

In paired daytime–nighttime casts on *AR75*, water-column eDNA seemed to shift upward at night (concentrations rose between roughly 15 and 90 m relative to 105–180 m), consistent with the upward leg of diel vertical migration (**Fig. S8**). In near-bottom samples from *PS2204*, distance above the seafloor was a poor predictor of total eDNA concentration (**Fig. S9**), suggesting that benthic boundary-layer effects do not dominate the signal in the way the depth attenuation does. Both observations are based on limited replication and should be treated as preliminary.

### 2. Comparative Assessment of Sampling Approaches

Since four expeditions used both rosette- and vehicle-mounted (ROV/HOV) Niskin bottle sampling, we tested whether the different approaches introduced systematic effects in the data. Vehicle operations may prolong the interval between water capture and shipboard filtration relative to rosette operations. A delay of several hours between collection, filtration, and stabilization could potentially lead to the degradation of some eDNA material ^51^

To minimize confounding, approach comparisons were restricted to expeditions that used both sampling approaches within identical environmental settings: pelagic samples from AL2019 (HOV n = 8 vs. rosette n = 43), demersal samples from PS2204 (ROV n = 57 vs. rosette n = 48), and demersal samples from FKt230417 (ROV n = 40 vs. rosette n = 6) (Fig. S3). While NF2202 vehicle data are illustrated, no formal comparison was conducted for this expedition because all vehicle samples were demersal while all rosette samples were pelagic.

Within each environmental stratum, log_10_-transformed eDNA concentration was regressed against log_10_ depth along with a sampling approach indicator variable. Because vehicle sample volumes are generally smaller than those of rosettes, we implemented two regression model formulations: (1) eDNA concentration alone, and (2) eDNA concentration with log_10_ filtered volume included as a covariate. Furthermore, attenuation exponents (b) were re-estimated for each productivity regime, with and without vehicle-collected samples, using Gamma log-link GLMs and applying regime-specific intercepts and expedition-clustered confidence intervals where applicable.

Concentration data from vehicle-mounted and CTD-rosette Niskin samples show no significant difference in *AL2019*, the one expedition in which the two approaches filtered overlapping water volumes (conc effect = 0.147, p = 0.252; conc_vadjusted effect = -0.073, p = 0.704). The two expeditions that have no volume overlap at all do show significant contrasts for concentration, but their effects have opposite directions and become non-significant after adjusting for volume (*PS2204* conc effect = -0.314, p < 0.0001; conc_vadjusted effect = -0.409, p = 0.054. *FKt230417* conc effect = 0.935, p = 0.002; conc_vadjusted effect = -3.447, p = 0.153). Further, removing all vehicle samples from the dataset shifts the depth-attenuation exponent *b* by at most 0.042 across all productivity regimes, which is well within the confidence intervals. Together, these analyses indicate that any residual differences between sampling approaches cannot be separated from differences in filtered volume, and that including vehicle-collected samples does not introduce bias in the depth-attenuation relationships that underpin this study.

### 3. Target eDNA measurements

Total eDNA is a heterogeneous mixture of DNA from diverse sources. Researchers studying eDNA are typically interested in targeting only a fraction of this diversity. To quantify the abundance of target eDNA relative to the total, we applied a new qPCR assay targeting a fragment of the ribosomal *28S* gene to the *FKt230417* sample set.

#### 3.1. Primers

This assay employed the primers developed by McCartin et al. ^3^: 5′-CGTGAAACCGYTRRAAGGG-3′ and 5′-TTGGTCCGTGTTTCAAGACG-3′. These primers were originally designed to selectively amplify a ∼388 bp *28S* region of anthozoan corals (sufficiently variable to distinguish genera and, in some taxa, species). Still, empirical results have demonstrated that they amplify across Cnidaria, Porifera, and Ctenophora ^3–6^. Primers were synthesized by Eurofins Genomics (Louisville, KY, USA), purified via standard desalting, and normalized to 100 M in TE buffer (pH 8.0). For experimental use, primer stocks were diluted to 10 µM using molecular-grade water.

Optimal primer concentrations were determined by comparing 300, 500, and 1000 nM concentrations on the generated standard curve template samples (see below). At 300 nM, R^2^ value was 0.977 with a doubling efficiency of 87%. At 500 nM, R^2^ value was 0.987 with a doubling efficiency of 83%. At 1000 nM, R^2^ value was 0.993 with a doubling efficiency of 88%. Based on the R^2^ value and doubling efficiency, a primer concentration of 1000 nM was used for subsequent qPCR assays.

#### 3.2. Sensitivity and quantification limits

Standard curves for sensitivity and quantification limit tests were synthesized as follows: The *28S* fragment of *Paracalyptrophora carinata* was amplified and cloned into a plasmid. Purified genomic DNA from *P. carinata* was amplified using the Taq DNA Polymerase with Standard Taq Buffer (New England Biolabs). The GeneJet gel extraction Kit (ThermoFisher Scientific) was used to isolate the target band, which was then ligated into a pGEM-T Vector (Promega, Madison, WI, USA) containing an ampicillin resistance gene. Plasmids were transformed into competent *E. coli* cells via heat shock at 42 °C for 35 seconds, followed by 2 minutes of cooling on ice. Cells were incubated at 37 °C in nutrient broth, plated on nutrient agar with ampicillin, and grown overnight. Success was confirmed by PCR amplification of the insertion using T7- and SP6-promoter-specific primers. Amplicons were visualized on a 1% TBE agarose gel stained with GelRed (Biotium, Fremont, CA, USA) and Sanger-sequenced to verify the target sequence. Final plasmids were purified using the GeneJET Plasmid Miniprep kit (ThermoFisher Scientific). DNA concentrations were determined using the Qubit 1X Broad Range dsDNA assay. Copy numbers were calculated based on the ratio of the amplicon mass (kDa) to the combined mass of the plasmid and insert. A standard curve was generated using a ten-fold serial dilution (ten standards starting with 1.0 x 10^10^ to 1 copy µL^-1^) of the purified plasmid vector.

Assay sensitivity and quantification limits were determined following Klymus et al. ^7^. The specific run used to define the LOQ/LOD yielded a slope of -3.364, an intercept of 35.113, and an R^2^ of 0.991. The limit of quantification (LOQ; coefficient of variation < 0.35) was determined to be 52 copies per reaction, and the modeled limit of detection (LOD) was 10 copies per reaction.

#### 3.3. qPCR Methodology

Quantitative PCR was performed in 20 µL reactions consisting of 10 µL of Quantabio PerfecTa SYBR Green Fastmix (Beverly, MA, USA), 1000 nM of each primer, 1 µL of bovine serum albumin (ThermoFisher Scientific), and 5 µL of 5x diluted template (5x diluted template to yield 1 copy rxn^-1^). All reactions were performed on a Qiagen Rotor-Gene 6000 Thermal Cycler using either 72- or 36-well rings. For 72-well configurations, clear Qiagen 0.1 mL four-strip tubes were utilized; for 36-well configurations, 0.2 mL single tubes designed for the RotorGene system were employed. Thermal cycling conditions were set as follows: initial denaturation at 95 °C for 10 min, followed by 40 cycles of 95 °C for 15 s, 70 °C for 15 s, and 72 °C for 30 s. Each run concluded with a melting curve analysis from 72 to 95°C. The Ct threshold was determined using the auto-threshold setting in the RotorGene software. Absolute concentrations (copies µL^-1^) were calculated by fitting Ct values to the run-specific standard curve. Across six qPCR runs, the average R^2^ value was 0.998 ± 0.001, and the doubling efficiency was 86 ± 1.6%.

#### 3.4. Contamination Controls and Field Monitoring

Contamination controls were implemented throughout the workflow, as described by McCartin et al. ^8^. Field contamination was monitored through sampling controls consisting of filtering ultrapure water generated onboard the *R/V Falkor (too)* using a Milli-Q Direct 8 after each sampling deployment. In the laboratory, these were processed in parallel with qPCR negative controls. Of the 7 sampling control samples (**Table S7**), all showed little to no amplification. When amplification occurred (3 of the 7), the measured concentration was an order of magnitude lower than that from any field sample.

### 4. Estimating filtration volume and extract concentration requirements for rank-n eDNA ASV detection

We developed a quantitative framework to estimate the seawater filtration volume required to detect the *n*-th most abundant amplicon sequence variant (ASV) in metabarcoding surveys, given measured eDNA concentration profiles and the downstream molecular workflow. The framework links three observational and modeling components: (i) a normalized power-law model of total eDNA concentration with depth fit per expedition, (ii) a Zipf-distributed rank-abundance model for the amplifiable target community, and (iii) an empirically-calibrated conversion between target eDNA mass and PCR copy number.

#### 4.1. Depth-dependent eDNA concentration model

Total eDNA concentration (*ρ_t_*), measured in nanograms per liter (ng L^-1^), was modeled as a normalized power law of depth:

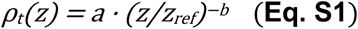

Where *z* is depth (m), *z*_ref_ is the assumed depth of the eDNA concentration maximum for that expedition (m), *a* is the power-law intercept, and *b* is the power-law attenuation exponent. Uncertainty in (log *a*, *b*) was propagated by drawing *n*_sims_ = 5000 joint samples from the bivariate normal distribution with mean (log *â*, *bR*) and covariance Σ. When the upstream fit provided a full variance–covariance matrix, Σ was used directly; when only marginal 95% confidence intervals were available, Σ was constructed under an independence assumption with marginal standard errors derived as (CI_upper_ − CI_lower_) / (2 × 1.96) for log(*a*) and *b*.

#### 4.2. Rank abundance

Following the empirical characterization and modeling of total eDNA yield declines with depth, we sought to parameterize the distribution of diversity within the target eDNA pool enriched by the McCartin et al. ^3^ *28S* primers (**Table S8)** to inform ASV-specific detection probabilities. We modeled the rank– abundance distribution of ASVs as a truncated Zipf distribution. We estimated the Zipf exponent *s* empirically from metabarcoding of *28S* amplicons generated by McCartin et al. ^5^ from *FKt230417* samples (Fig. S10).

Under a Zipf model, the fractional contribution of the *n*-th ranked ASV to the *28S*-amplifiable community is:

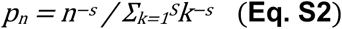

where *p_n_* is the fractional contribution of the *n*-th ranked ASV (dimensionless), *n* is the rank of the ASV, *S* is the maximum number of distinct *28S* ASVs expected per sample, and *s* is the Zipf exponent. Higher values of *s* indicate a community more heavily dominated by a small number of ASVs, and *s* therefore describes the steepness of the dominance–diversity curve and, equivalently, the relative evenness of the community: low *s* (≲ 1) indicates an even community in which rare ASVs collectively contribute a substantial fraction of total eDNA, whereas high *s* (≳ 3) indicates a community in which rare ASVs contribute negligible fractions and would require correspondingly large filtration volumes to detect.

#### 4.3. Quality control and control-sample exclusion

For each sample, we required at least 1000 total *28S* reads and at least 10 non-zero ASVs to be eligible for Zipf fitting. Control samples were excluded from all rank-abundance, richness, and downstream summaries. The QC outcome and per-sample *s* estimates for every QC-passing sample are reported in **Table S9**.

#### 3.4. Empirical estimation of community size S

Rather than fixing *S* to a single global value, we set *S* per sample based on observed ASV richness (the number of ASVs with at least one read). For sensitivity analysis, we additionally computed the bias-corrected Chao1 estimator ^9^ of true (including unobserved) richness:

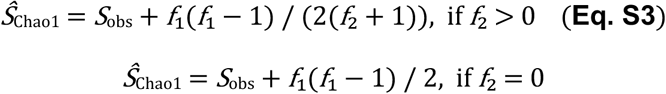

where *f*₁ and *f*₂ are the counts of singleton and doubleton ASVs, Per-sample *S*_obs_, *f*₁, *f*₂, and Chao1 estimates are reported in **Table S9**. Observed ASV richness and Chao1 were consistent and nearly identical.

#### 4.5. Discrete maximum likelihood estimation of s

We estimated *s* per sample by discrete maximum likelihood. Under the multinomial model with cell probabilities given by **Eq. S2**, the log likelihood for observed read counts *c_n_* at ranks *n* = 1, …, *N* (with *N* ≤ *S*) is, up to a constant in *s*,

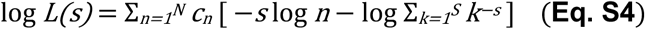

The MLE was obtained by numerical minimization of −log *L(s)* over *s* ∈ [0.1, 8] using the stats::optimize R function. The likelihood-based estimator is preferred over log–log linear regression because it correctly weights ranks by their observed read counts and incorporates the truncation *S* through the normalizing sum. For comparison, we computed unweighted log–log OLS and read-count-weighted OLS for every sample; these are reported alongside the MLE in **Table S9**. The expedition-level median of per-sample MLEs for samples collected deeper than 200 m was used as the *s* parameter in downstream calculations **Table S10**. Per-expedition-by-depth-zone summaries (epipelagic 0–200 m, mesopelagic 200–1,000 m, bathypelagic > 1,000 m) are summarized in **Table S11**.

#### 4.6. 28S mass to copy conversion

The *28S* copy concentration of the n-th ranked ASV (*ρ_rank_*) was derived from total eDNA via:

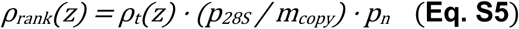

Where *ρ_rank_*(*z*) is the *28S* copy concentration of the *n*-th ranked ASV (copies L^-1^), *p_28S_* is the *28S* mass fraction of total eDNA, *m_copy_* is the mass per amplicon molecule (≈ 388 bp), which is 4.19 × 10^-10^ ng copy^-1^ (assuming an average molecular weight of a base pair in dsDNA of 650 g mol^-1^ bp^-1^), and *ρ_t_*(*z*), and *p_n_* are as defined above.

Where paired bulk eDNA (Qubit) and *28S* qPCR measurements were available for the expedition, *p_28S_* was empirically calibrated as the mean of per-sample target-to-total ratios *p_28S_* = (qPCR copies L^-1^ *· m_copy_*) / (total ng L^-1^). For the *FKt230417* expedition, this calibration yielded *p_28S_* = 2.5 × 10^-7^ (dimensionless).

#### 4.7. Volume requirement at depth

The filtration volume *V_rank_(z)* required to deliver at least *ρ_min_* copies of the n-th ranked ASV into each PCR reaction is:

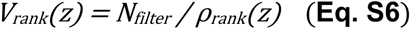

Where *V_rank_*(*z*) is the filtration volume required (L), *N_filter_* is the filter-side copy target (copies), and *ρ_rank_*(*z*) is as defined above. The filter-side copy target *N_filter_* accounts for extraction efficiency *E* and the dilution introduced by loading only *V_template_* µL of template from a *V_elution_* µL extract:

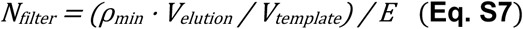

Where *ρ_min_* is the minimum copy target per PCR reaction (copies/well, set to 20 copies rxn^-1^), *V_elution_* is the extract elution volume (set to 80 µL), *V_template_* is the template volume loaded into the PCR reaction (set to 5 µL), and *E* is the extraction efficiency (dimensionless, set to 0.80). We set *ρ_min_* = 20 copies rxn^-1^ as the conservative threshold for consistent amplification and detection by qPCR (equivalent to 2x qPCR LOD; ^10,11^), *E* = 0.80 based on published recoveries for our extraction kit, and *V_template_* = 5 µL drawn from a *V_elution_* = 80 µL extract.

#### 4.8. Minimum extract concentration

The minimum bulk extract concentration (*ρ_ng/µL_^min^*) that guarantees rank-*n* detection follows from rearranging the PCR template chain:

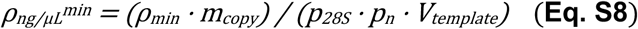

Where *ρ_ng/µL_^min^* is the Minimum total extract concentration (ng µL^-1^), and *ρ_min_*, *m_copy_*, *p_28S_*, *p_n_*, and *V_template_* are as defined above.

#### 4.9. Sensitivity analyses

We performed two complementary sensitivity analyses to characterize how the volume and concentration requirements respond to plausible variation in the molecular parameters. First, we evaluated **Eq. S8** analytically across all factorial combinations of five parameters (*V_elution_* ∈ {20, 40, 60, 80, 100} µL; *V_template_* ∈ {1, 2, 5, 10} µL; *ρ_min_* spanning 3 to 40 copies rxn^-1^; *s* spanning 1.0 to 3.0; *n* spanning 10 to 60) (**Figure S6)**. Because **Eq. S8** is closed-form (generates an exact answer), no Monte Carlo propagation was needed, and the full factorial of 1600 combinations was evaluated. Second, we re-evaluated the depth-dependent volume requirement *V_rank_*(*z*) across the same parameter grid using Monte Carlo propagation of the power-law uncertainty, with cached (*a*, *b*) draws reused across scenarios for computational efficiency (**Fig. S7)**. Outputs include median and 5–95% / 25–75% uncertainty bands at depths spanning 200–4000 m.

## SUPPORTING INFORMATION FIGURES

**Figure S1.**
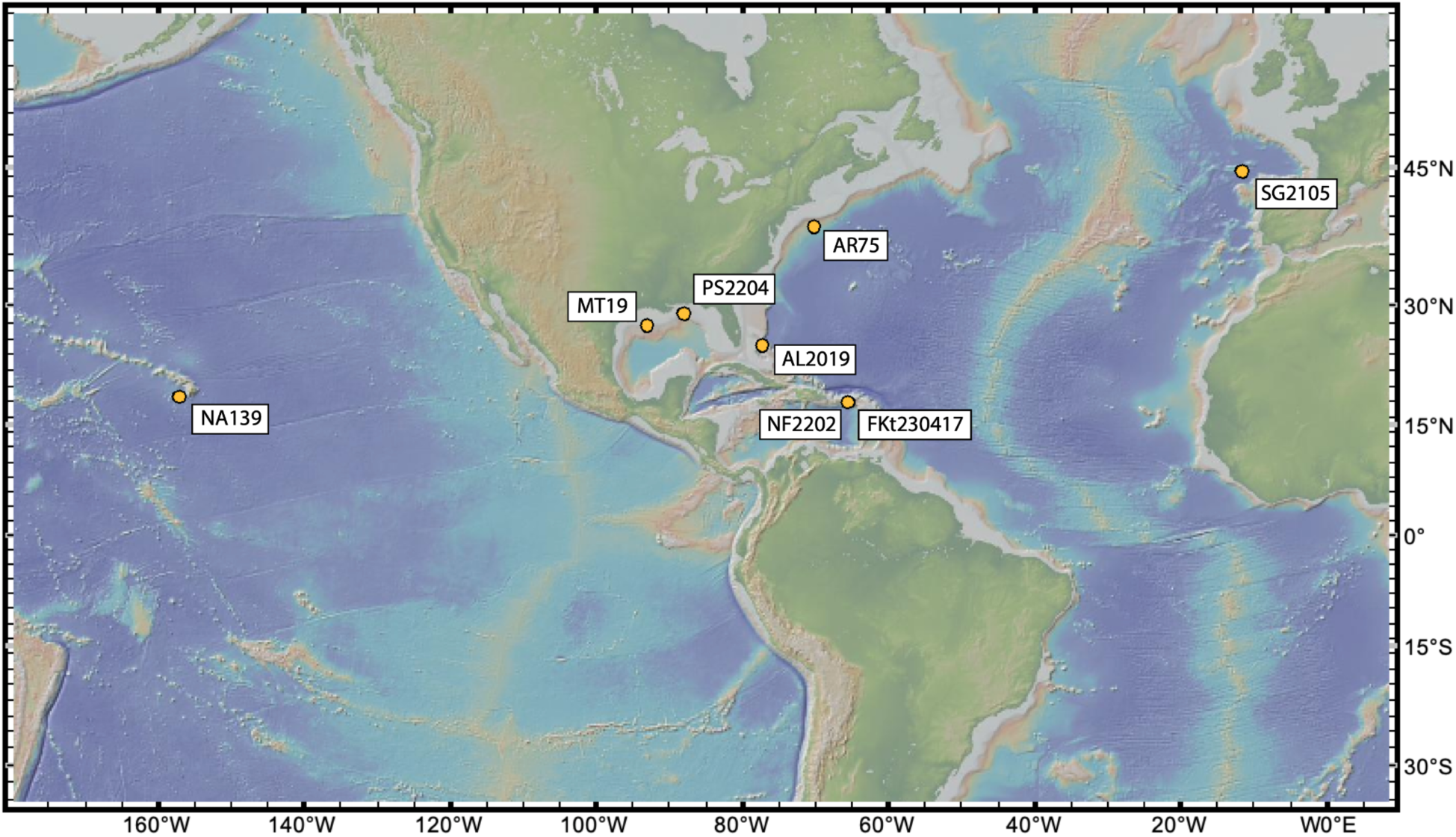
Map of oceanographic expeditions where eDNA was collected for this study.

**Figure S2.**
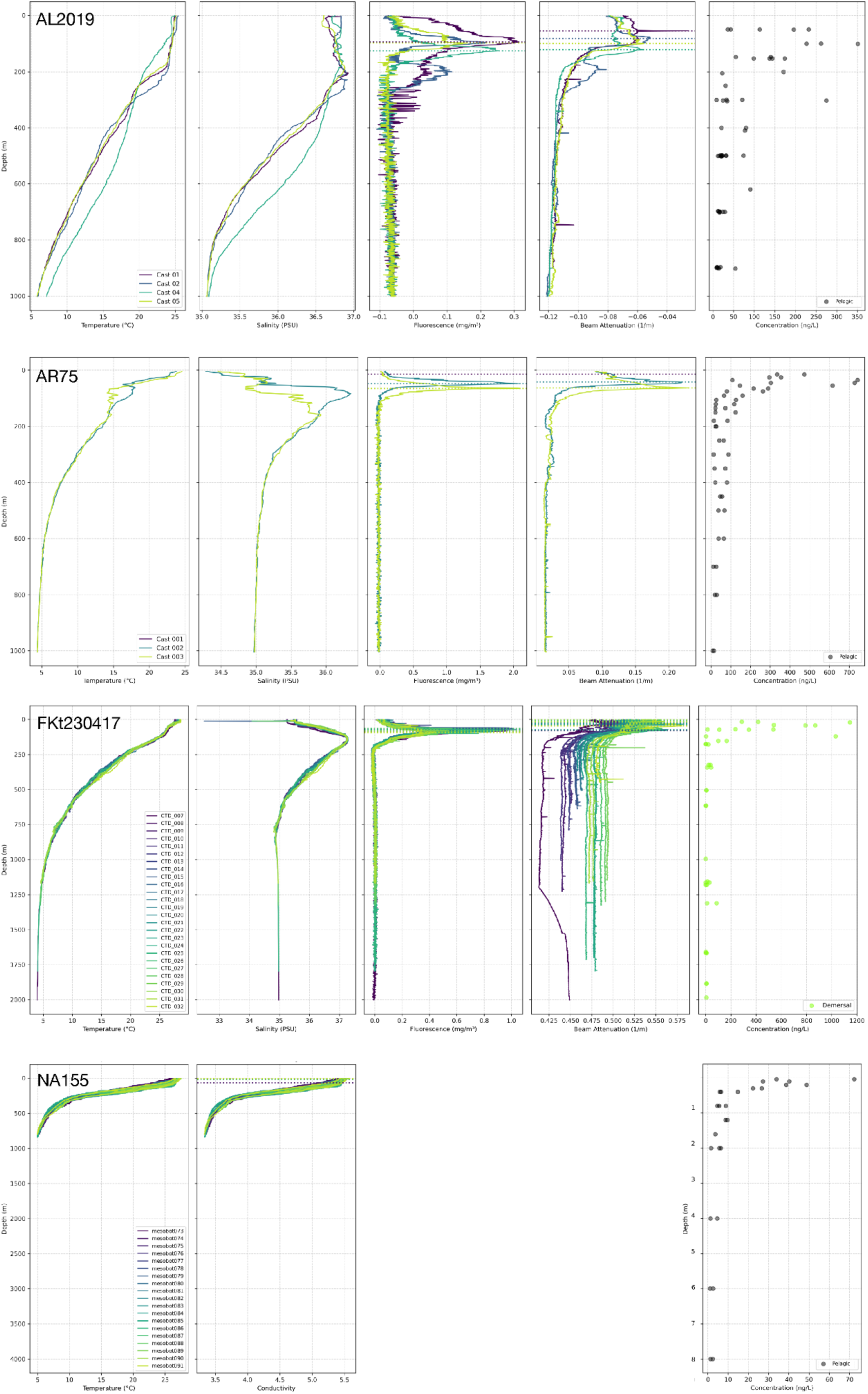
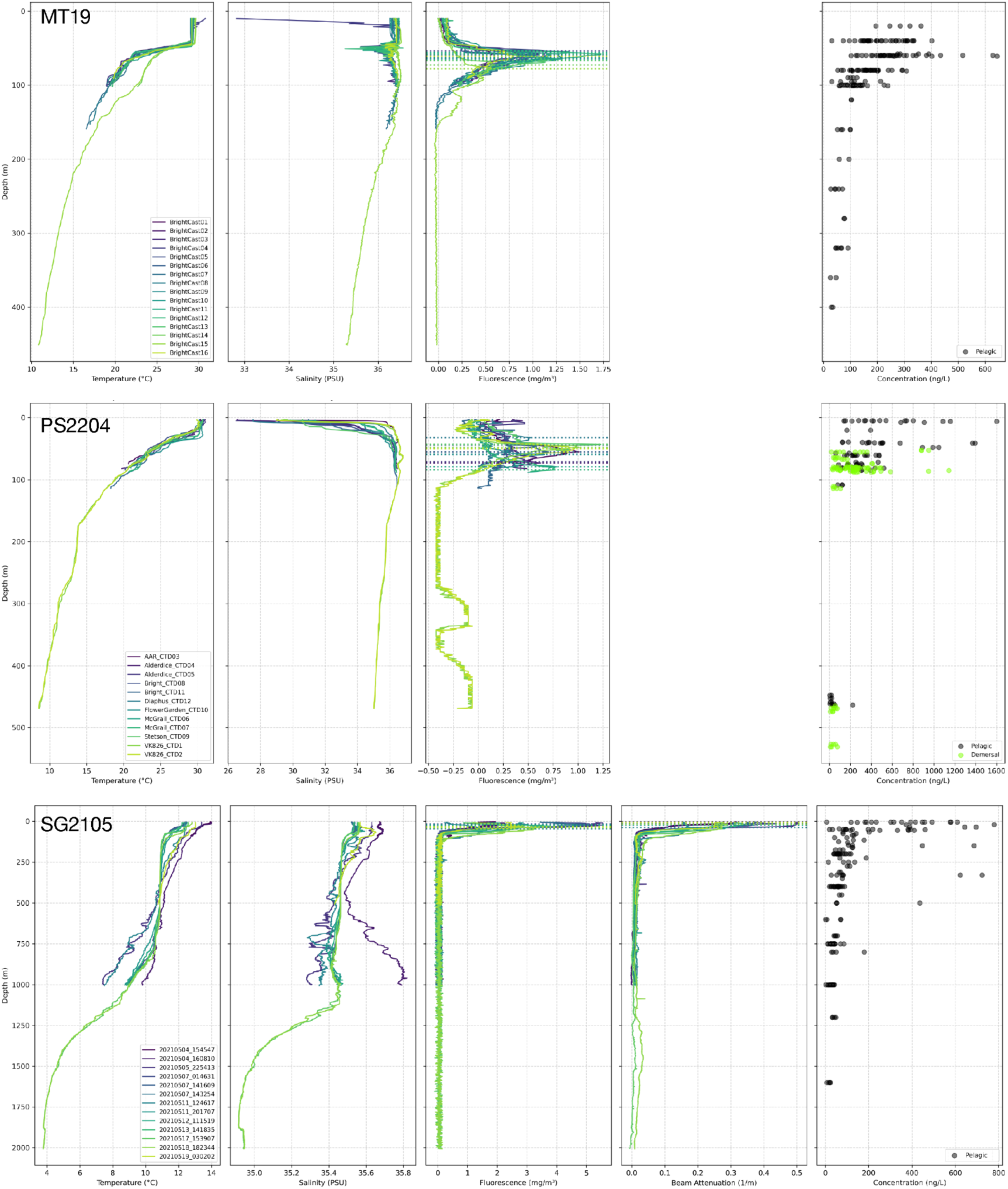
Vertical profiles of physical, biological, and chemical parameters across eight research expeditions. Each row presents data from a single expedition. Columns from left to right show: temperature (°C), salinity (PSU/conductivity), fluorescence (mg/m³), beam attenuation (1/m), and pelagic or demersal total eDNA concentration measurements (ng/L) plotted against depth (m). Individual casts/deployments are distinguished by color, and dotted horizontal lines in the fluorescence and beam attenuation panels indicate the depth of the deep chlorophyll maximum (DCM) and the beam attenuation maximum for each cast/deployment. Total eDNA concentration measurements are categorized as pelagic (gray) or demersal (green), as indicated in each panel’s legend. Note differences in depth ranges and x-axis scales among expeditions. Fluorescence and beam attenuation profiles were not available for NA155. Beam attenuation profiles were not available for MT19 or PS2204.

**Figure S3.**
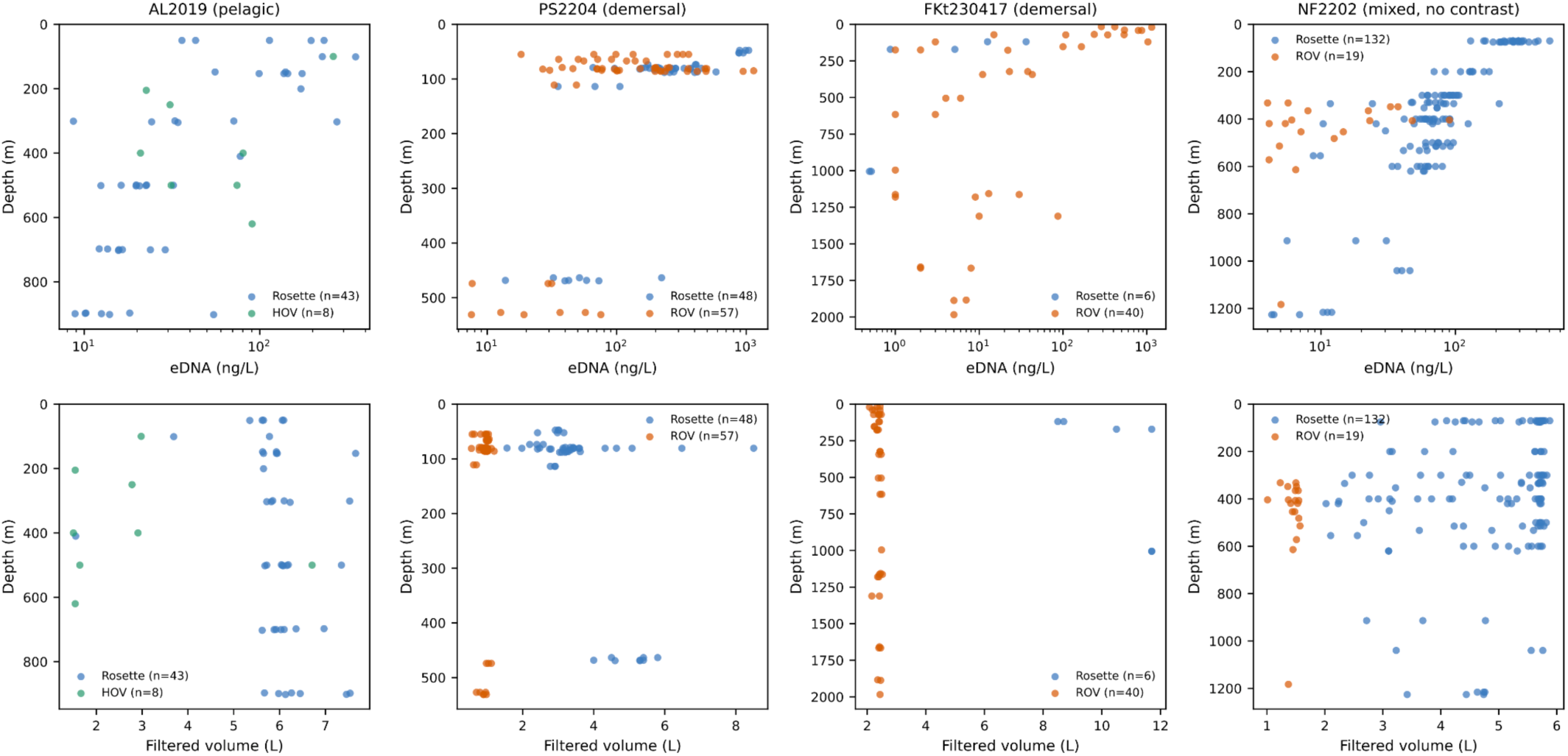
Comparison of rosette-mounted and vehicle-mounted (ROV or HOV) Niskin sampling. Total eDNA concentration is plotted against depth for the four expeditions that sampled with both approaches, with points colored by platform. Contrasts controlling for depth within a single environment are not significant in pelagic samples and differ in sign between demersal expeditions, and the fitted depth-attenuation exponents change by less than 0.05 when all vehicle-mounted samples are excluded.

**Figure S4.**
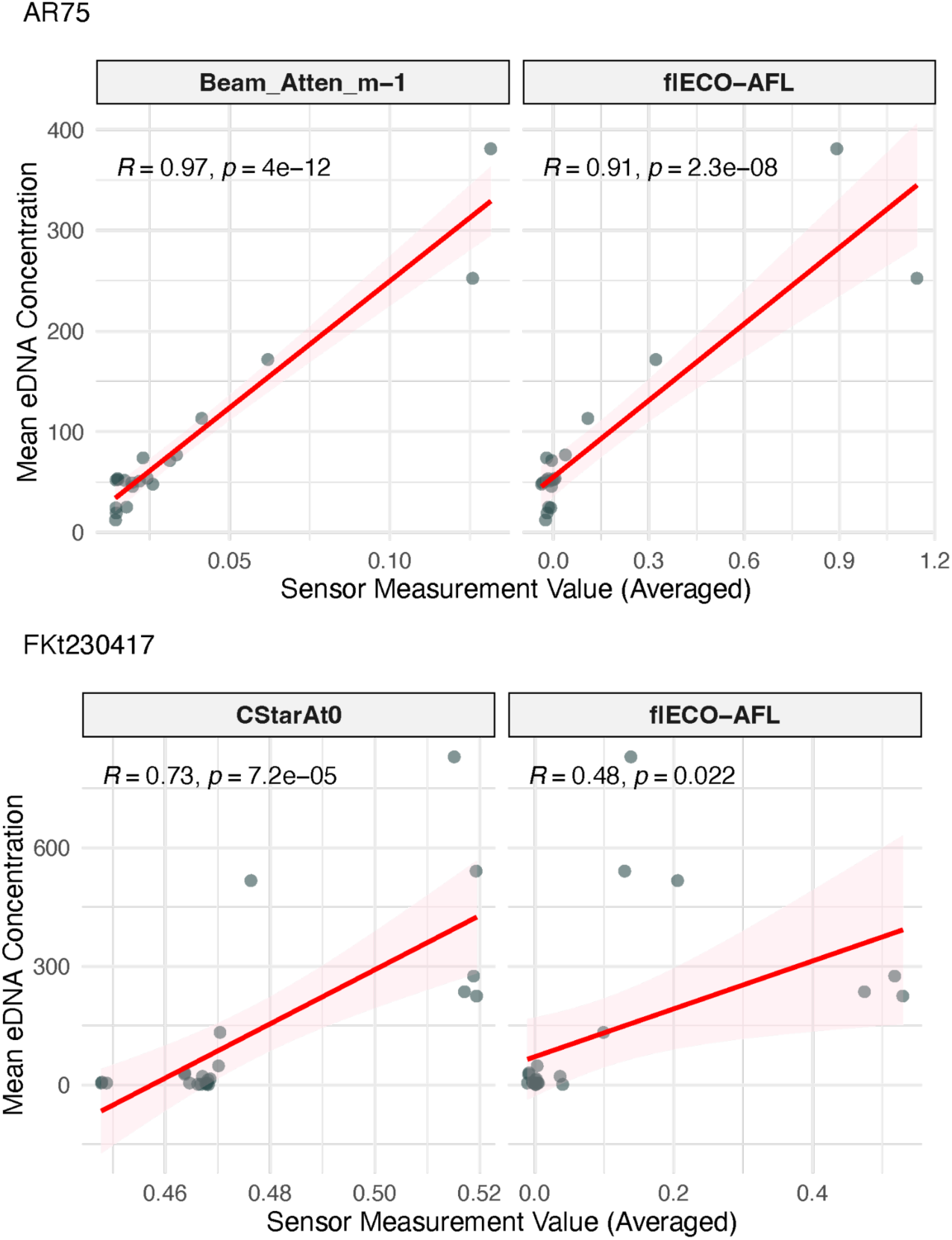
Relationship between eDNA yields and in situ CTD sensor measurements. Representative data from expeditions AR75 (top panels) and FKt230417 (bottom panels) were averaged by depth. Panels show the correlation between eDNA yield and fluorescence (left) and between eDNA yield and beam attenuation (right). The red line indicates the linear regression fit (with pink shading denoting the 95% confidence interval). Statistical significance and effect size are quantified via Pearson’s R and associated p-values in the top left of each panel.

**Figure S5.**
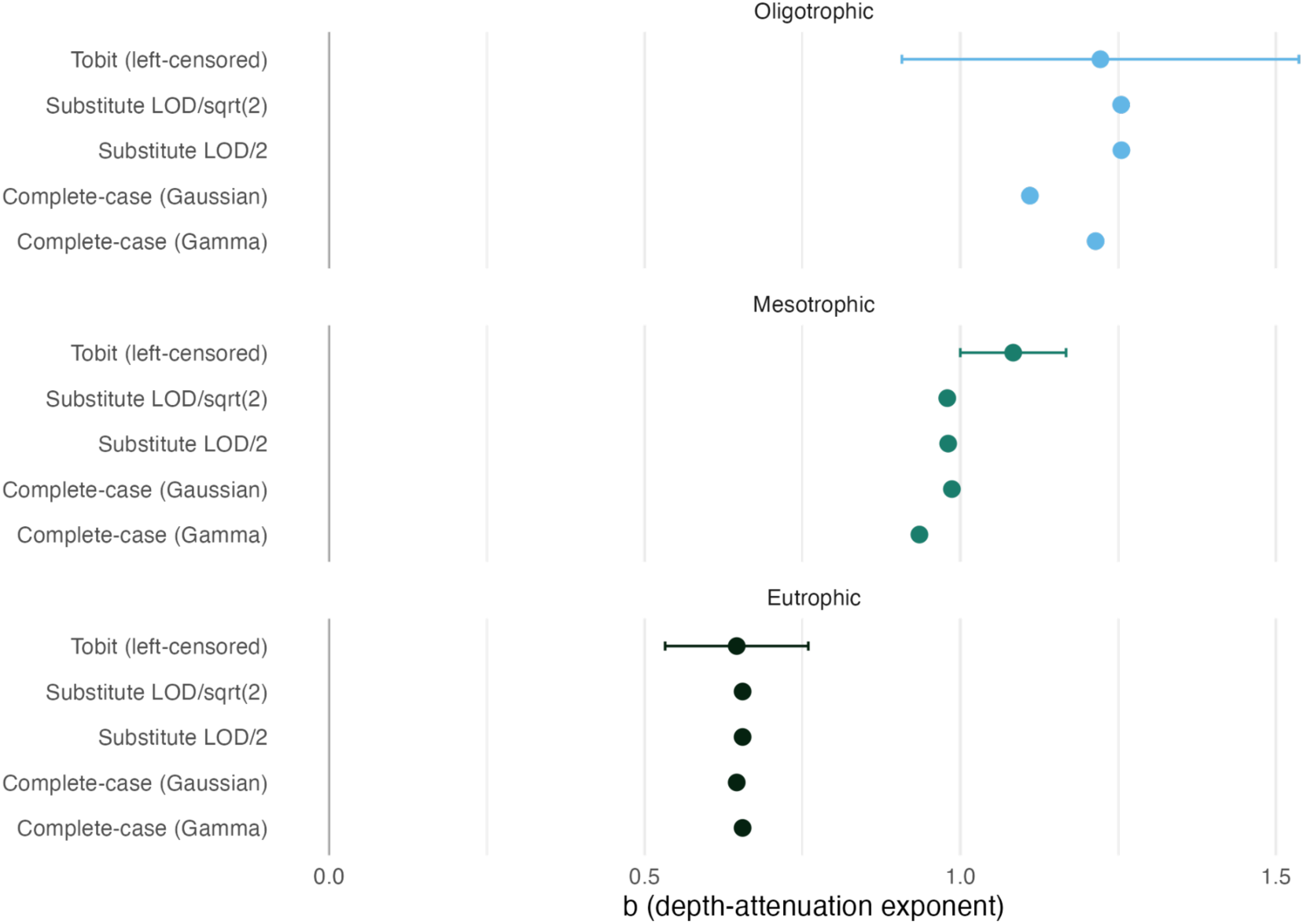
Depth-attenuation exponent b under each treatment of samples at or below the assay quantification limit, by productivity regime. Points show b from the complete-case Gamma-log GLM, complete-case Gaussian regression on log-concentration, substitution at LOQs/2 and LOQs/√2, and left-censored (Tobit) regression; horizontal bars give the Tobit 95% confidence interval. Non-detect counts and their depth distribution are summarized alongside.

**Figure S6.**
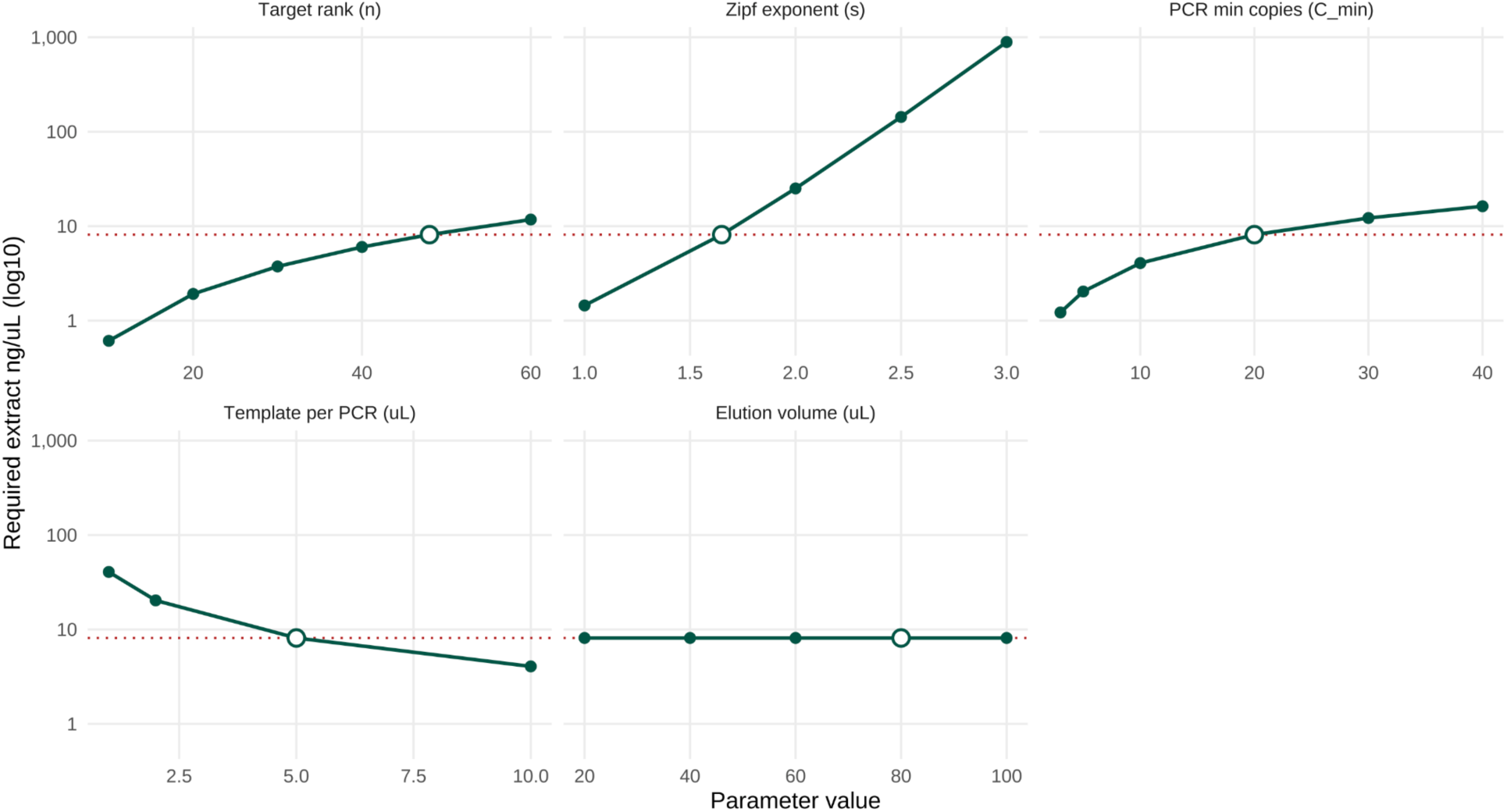
Sensitivity of the minimum extract concentration *ρ*_ng/µL_^min^ required to deliver a target number of 28S amplicon copies per PCR reaction for the n-th ranked ASV in expedition FKt230417. *ρ*_ng/µL_^min^ is computed from the expression in **Eq. S8** of the Supporting Information, which links the rank-abundance Zipf model to the loaded PCR template, and is independent of depth. The five panels sweep one parameter at a time (target rank n, Zipf exponent *s*, minimum PCR copy threshold *ρ*_min_, template volume per PCR *V*_template_, and elution volume *V*_elution_) across the same grids used in the depth-explicit volume sensitivity (**Fig. S7**); the open circle on each panel marks the baseline value, and the red dotted horizontal reference line marks the baseline *ρ*_ng/µL_^min^ = 8.1 ng µL⁻¹. The y-axis is on a log₁₀ scale. Because **Eq. S8** is analytic, no Monte Carlo propagation is needed, and the curves shown are exact under the model. Note that *V*_elution_ does not affect *ρ*_ng/µL_^min^ because the dilution introduced by a larger elution is exactly canceled by the smaller fraction loaded per PCR (*V*_template_ / *V*_elution_); the panel is included to illustrate this. Of the four parameters that do affect *ρ*_ng/µL_^min^, the Zipf exponent *s* exerts the largest influence (an approximately three-order-of-magnitude change across the 1.0–3.0 range), reflecting the strong effect of community evenness on the per-rank fractional contribution.

**Figure S7.**
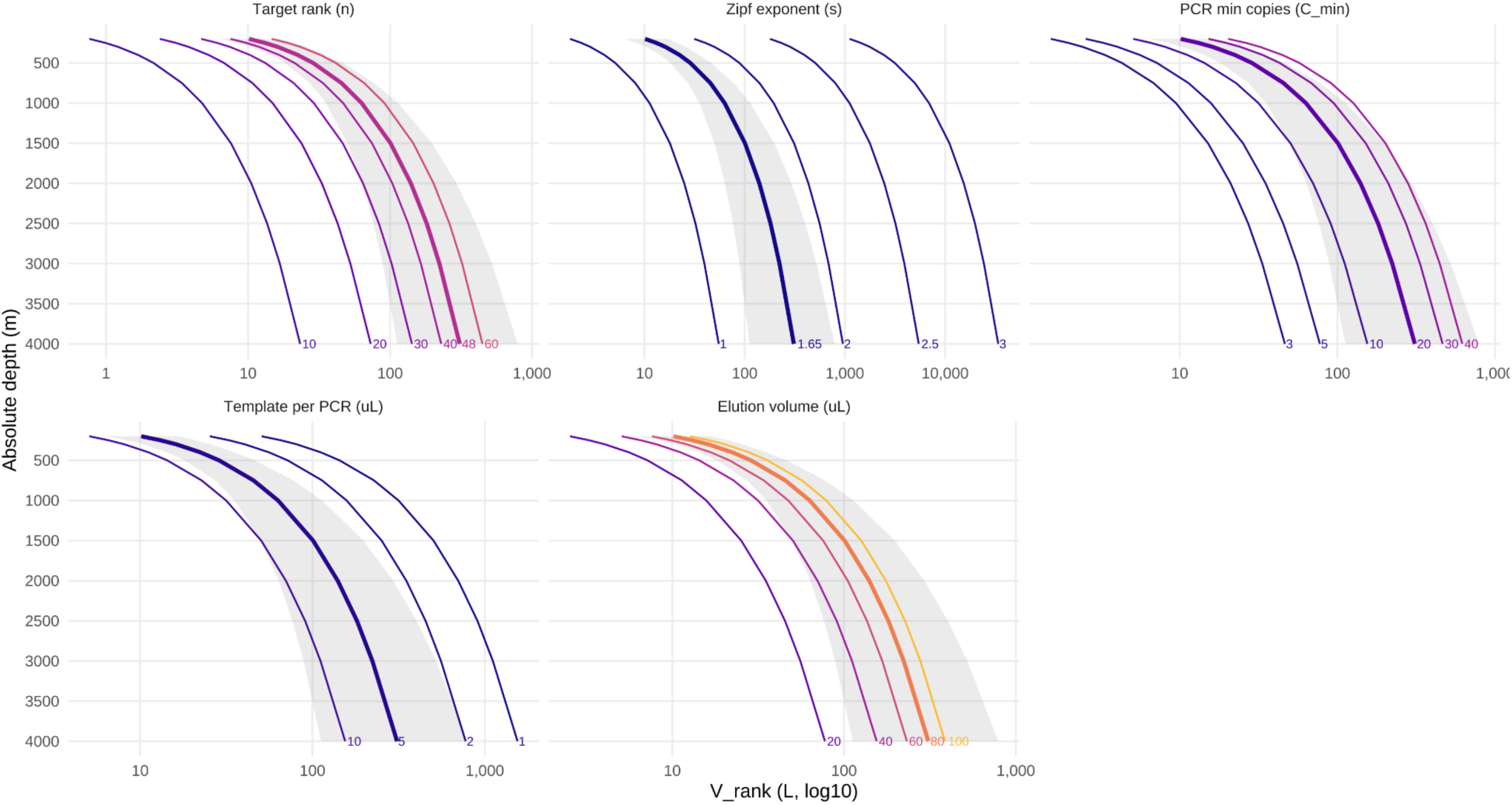
Sensitivity of the seawater filtration volume *V*_rank_(*z*) required to detect a rank-n target ASV by qPCR or amplicon sequencing, evaluated for expedition *FKt230417* (rawXexpedition model). Predictions are derived from **Eq. S6** of the Supporting Information, which combines the depth-attenuation power-law for total eDNA, a Zipf rank-abundance model for the 28S-amplifiable community, and a copy-number target on the loaded PCR template. Each panel sweeps one parameter at a time around its baseline value (thick colored line), while the remaining four are held fixed; the grey ribbon shows the 5–95% Monte Carlo uncertainty band at baseline, propagated from the variance–covariance matrix of the fitted power-law parameters (5,000 joint draws). Baseline values: target rank n = 48 (i.e., the 95th-percentile-rare ASV in a community of S = 50 distinct 28S ASVs), Zipf exponent *s* = 1.65 (empirical median across FKt230417 samples below 200 m), *ρ*_min_ = 20 copies per PCR well, *V*_template_ = 5 µL, *V*_elution_ = 80 µL, extraction efficiency *E* = 0.80, p_28S = 2.5 × 10⁻⁷ (empirically estimated from paired Qubit and 28S qPCR measurements on the same samples), and *Z_ref_* = 25 m. Parameter values are labeled at the deepest plotted depth; volume axes are on a log₁₀ scale. Note that *V*_elution_ does not appear in **Eq. S6** (increasing the elution volume dilutes the extract but loads the same fraction per PCR) and is shown here only to illustrate cancellation of dilution and concentration effects across the range tested. Among the parameters that scale *V*_rank_, the Zipf exponent *s* exerts the largest leverage (≈ 100-fold change across the 1–3 range), followed by target rank n (≈ 30-fold across 10–60), *ρ*_min_ (≈ 13-fold across 3–40), and *V*_template_ (≈ 10-fold across 1–10 µL).

**Figure S8.**
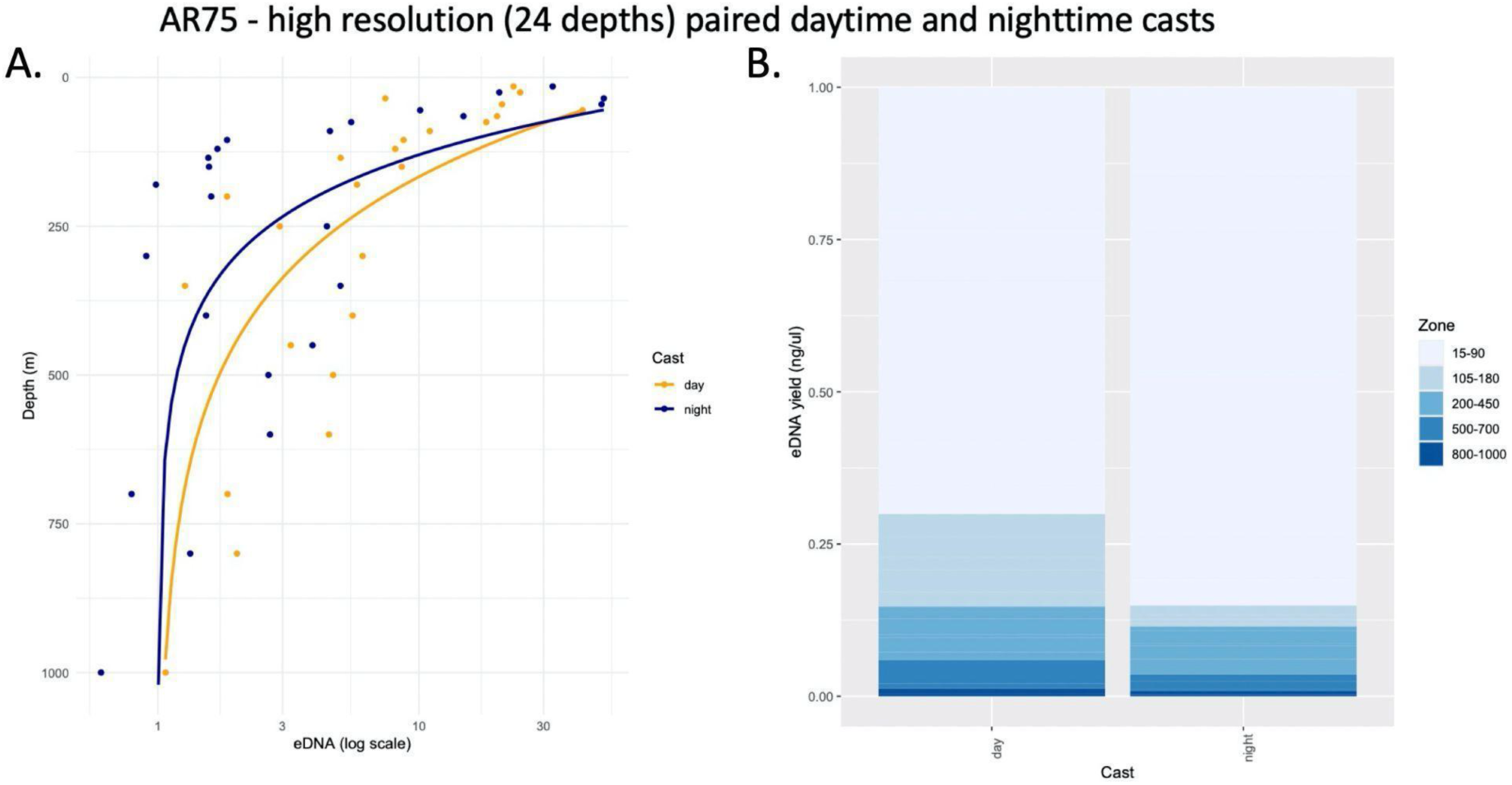
In the *AR75* expedition, back-to-back high-resolution vertical resolution (24 depths) daytime and nighttime eDNA collections were obtained from CTD casts. A) Depth vs. eDNA yield for daytime (yellow) and nighttime (blue). B) Proportion of eDNA in different depth zones for day (left) and night (right). At night, a proportionately greater amount of eDNA is in the 15-90 m zone.

**Figure S9.**
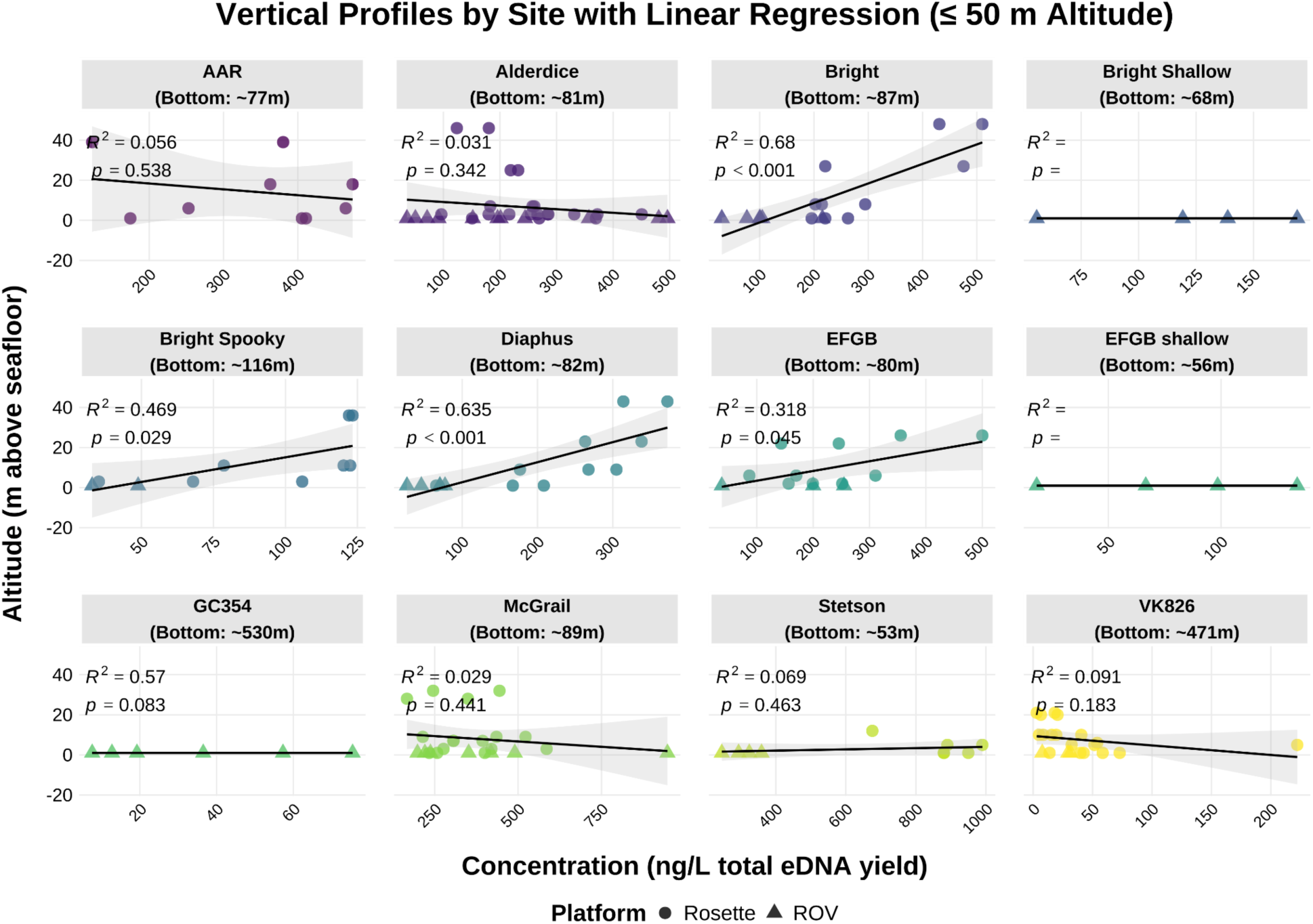
Vertical profiles of total eDNA concentration in the near-bottom boundary layer across sampling sites during expedition *PS2204*. Each panel shows total eDNA concentration (ng/L) plotted against altitude above the seafloor (m) for samples collected within 50 m of the bottom, with the bottom depth estimated as the mean of the sample depth and the mean altitude over the site. Points represent individual samples collected using Niskin bottles on an ROV or CTD Rosette; the solid black line and shaded band show the per-site ordinary least-squares regression fit with 95% confidence interval. R² and p-values are reported for each site. The strength and direction of the concentration–altitude relationship varied among sites, suggesting that vertical eDNA gradients in the near-bottom boundary layer are spatially heterogeneous rather than following a consistent dilution pattern with increasing distance from the seafloor.

**Figure S10.**
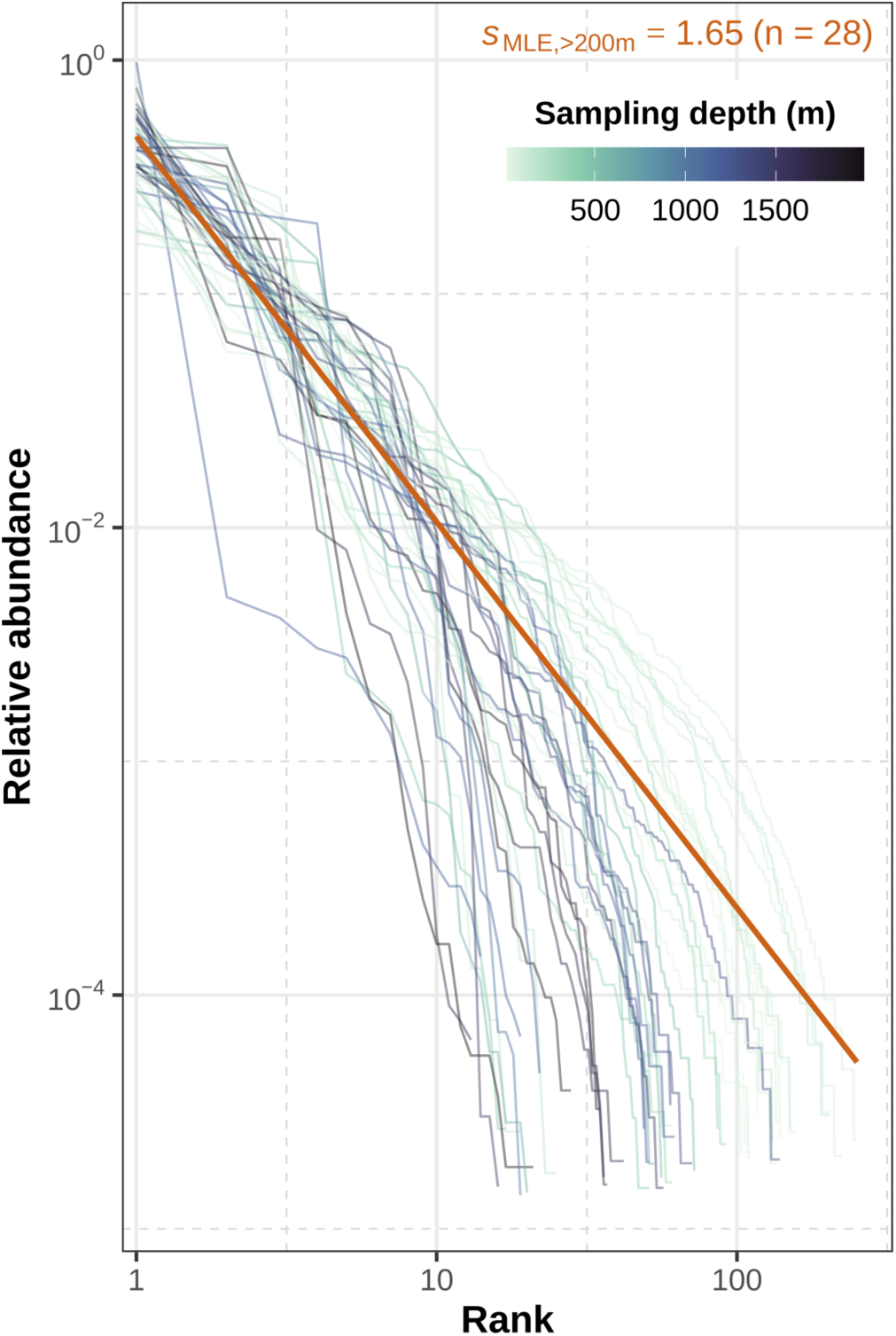
Rank–abundance distributions of 28S ASVs for *FKt230417* with fitted Zipf curve. Lines represent the empirical rank–abundance curves for each sample from *FKt230417* sequenced by McCartin et al. 2026. Lines are colored by sampling depth. For each sample, ASV read counts were ranked in descending order and converted to relative abundances. The orange line is the Zipf distribution evaluated using the median (below 200 m) of per-sample maximum-likelihood Zipf exponents (*s*_MLE_) and a truncation *S* set to the maximum per-sample observed ASV richness. Median *s*_MLE_ and sample size are indicated. Both axes are on a log₁₀ scale.

## SUPPORTING INFORMATION TABLES

**Table S1.** Per-sample eDNA quantifications and collection metadata for the 841 samples compiled from the eight oceanographic expeditions analyzed in this study, of which 756 were retained after the expedition-specific depth cutoffs and 729 after the further exclusion of samples at or below the assay limit of quantification. Variables include eDNA mass recovered per filter (ng), filtered seawater volume (L), sampling depth (m), the resulting concentration (ng L⁻¹), and metadata describing collection site, filter type, extraction method, expedition, sampling platform (Niskin rosette, ROV, or HOV), region, climate zone, productivity regime, collection date, and sample environment (pelagic or demersal).

**Table S2.** Per-sample eDNA quantifications and collection metadata for the samples at or below the Qubit assay quantification limit. The table mirrors the schema of Table S1 (collection metadata, filtered volume, depth) and adds the elution volume (µL) and the Qubit assay used, from which each sample’s limit of quantification in ng L⁻¹ is reconstructed for the censored-data reanalysis (Table S3, Fig. S5).

**Table S3**. Sensitivity of the depth-attenuation power law to the inclusion of samples at or below the assay quantification limit. Rows report, for each aggregation level (all samples, productivity regime, or individual expedition), the fitted exponent b under five treatments: complete-case Gamma-log GLM (the primary analysis), complete-case Gaussian regression on log-concentration, substitution of non-detects at LOQs/2 and at LOQs/√2, and left-censored (Tobit) maximum-likelihood regression with each non-detect censored at its sample-specific limit of quantification. Also reported are the Tobit 95% confidence interval, the numbers of detected and non-detected samples, and the shift in b between the censored and complete-case fits.

**Table S4.** Fitted parameters of the depth-attenuation power law (**Eq. 1**) for every model variant reported in this study, with rows corresponding to each combination of aggregation level (all samples, productivity regime, or individual expedition) and model type (raw or volume-weighted normalized-depth bin). Reported quantities are the back-transformed intercept *a* (ng L⁻¹), the attenuation exponent *b*, their 95% confidence intervals, the pseudo-*R*², and the marginal variances and covariance of (log *a*, *b*) used for Monte Carlo uncertainty propagation.

**Table S5.** Predicted seawater filtration volume *V*(*Z*) (L) as a function of depth (m) and productivity regime when one of three operational parameters, target extract concentration *ρ*t (ng µL⁻¹), elution volume *V*e (µL), or extraction efficiency *E*, is swept across its tested range while the other two are held at baseline (*ρ*t = 8.1 ng µL⁻¹, *V*e = 80 µL, *E* = 0.80). Each row reports the median *V*(*Z*) and the 5th, 25th, 75th, and 95th Monte Carlo percentiles. This table contains the underlying data for **Fig. 3**.

**Table S6.** Predicted seawater filtration volume *V*(*Z*) (L) across the full factorial of *ρ*t (0.5–20 ng µL⁻¹, 7 levels), *V*e (20–150 µL, 6 levels), and *E* (0.20–1.00, 5 levels) at depths from 200 to 4000 m and across the three productivity regimes. Each row reports the median *V*(*Z*) and the 5th, 25th, 75th, and 95th Monte Carlo percentiles. This factorial defines the ∼1500-fold operational envelope (0.012× to 18.5× the baseline).

**Table S7.** Paired total and 28S-target eDNA quantifications for *FKt230417* Niskin samples (Puerto Rico). For each sample, the table reports collection metadata (site, depth in m, filtered volume in L, filter type, filtration method), Qubit-quantified total eDNA (extract concentration in ng µL⁻¹, total filter mass in ng, and seawater concentration in ng L⁻¹), 28S-target eDNA from qPCR (copies per reaction, copies L⁻¹, and the derived target mass in ng L⁻¹ assuming 4.19 × 10⁻¹⁰ ng copy⁻¹), and the resulting target-to-total ratio. This table contains the underlying data for **Fig. 4**.

**Table S8.** Absolute abundances of 28S amplicon sequence variants (ASVs) in the FKt230417 Niskin samples, obtained by rescaling each ASV’s relative read abundance by the total 28S copy number measured by qPCR in the same extract. Columns report the sample identifier, expedition, depth (m), ASV identifier and taxonomic assignment, relative read abundance, and the resulting absolute copy concentration (copies L−¹). These data underlie the rank-abundance analyses in Tables S9 to S11.

**Table S9.** Per-sample fits of the truncated Zipf rank-abundance distribution (**Eq. S2**) to 28S metabarcoding ASV counts from QC-passing *FKt230417* samples sequenced by McCartin et al. 2026. For each sample, the table reports three estimators of the Zipf exponent *s* — discrete maximum likelihood (*s*MLE), unweighted log–log OLS, and read-count-weighted OLS — alongside observed and Chao1-corrected richness (with singleton and doubleton counts), total reads, sampling depth (m), and depth zone.

**Table S10.** Expedition-level summary statistics of the per-sample Zipf estimates for QC-passing, non-outlier samples collected deeper than 200 m for FKt230517. Reported quantities are the mean, median, SD, minimum, and maximum of each of the three Zipf estimators (MLE, unweighted OLS, weighted OLS) and of observed and Chao1 richness, together with the depth range spanned (m). The median *s* MLE reported here is the value used in **Eq. S2** for rank-abundance calculations.

**Table S11.** Depth-zone summary of the per-sample Zipf estimates, disaggregated as Epipelagic (0–200 m), Mesopelagic (200–1000 m), and Bathypelagic (>1000 m) for FKt230517. Same summary statistics as **Table S10**. This table llustrates how the dominance–evenness structure (Zipf exponent *s*) of the 28S-amplifiable community shifts with depth.

